# Optogenetic control of a horizontally acquired region in yeast prevent stuck fermentations

**DOI:** 10.1101/2024.07.09.602721

**Authors:** David Figueroa, Diego Ruiz, Nicolò Tellini, Matteo De Chiara, Eduardo I. Kessi-Pérez, Claudio Martínez, Gianni Liti, Amparo Querol, José M. Guillamón, Francisco Salinas

**Affiliations:** Laboratorio de Genómica Funcional, Instituto de Bioquímica y Microbiología, Facultad de Ciencias, Universidad Austral de Chile, Valdivia, Chile; ANID–Millennium Science Initiative–Millennium Institute for Integrative Biology (iBio), Santiago, Chile; Université Côte d’Azur, CNRS, INSERM, IRCAN, Nice, France; Centro de Estudios en Ciencia y Tecnología los Alimentos (CECTA), Universidad de Santiago de Chile (USACH), Santiago, Chile; Departamento de Ciencia y Tecnología de los Alimentos, Facultad Tecnológica, Universidad de Santiago de Chile (USACH), Santiago, Chile; Departamento de Biotecnología de los Alimentos, Instituto de Agroquímica y Tecnología de los Alimentos – Consejo Superior de Investigaciones Científicas (CSIC), Valencia, Spain

**Keywords:** Horizontal gene transfer, optogenetics, yeast, gene expression, fermentation

## Abstract

Nitrogen limitations in the grape must is the main cause of stuck fermentations during the winemaking process. In *Saccharomyces cerevisiae*, a genetic segment known as region A, which harbors 12 protein-coding genes, was acquired horizontally from a phylogenetically distant yeast species. This region is mainly present in the genome of wine yeast strains, carrying genes that have been associated with nitrogen utilization. Despite the putative importance of region A in yeast fermentation, its contribution to the fermentative process is largely unknown. In this work, we used a wine yeast strain to evaluate the contribution of region A to the fermentation process. To do this, we first sequenced the genome of the wine yeast strain known as ‘ALL’ using long-read sequencing and determined that region A is present in a single copy with two possible subtelomeric locations. We then implemented an optogenetic system in this wine yeast strain to precisely regulate the expression of each gene inside this region, generating a collection of 12 strains that allow for light- activated gene expression. To evaluate the role of these genes during fermentation, we assayed this collection using microculture and fermentation experiments in synthetic must with varying amounts of nitrogen concentration. Our results show that changes in gene expression for genes within this region can impact growth parameters and fermentation rate. We additionally found that the expression of various genes in region A is necessary to complete the fermentation process and prevent stuck fermentations under low nitrogen conditions. Altogether, our optogenetics-based approach demonstrates the importance of region A in completing fermentation under nitrogen-limited conditions.

**IMPORTANCE:** Stuck fermentations due to limited nitrogen availability in grape must represents one of the main problems in the winemaking industry. Nitrogen limitation in grape musts reduce yeast biomass and fermentation rate, resulting in incomplete fermentations with high levels of residual sugar, undesired by-products, and microbiological instability. Here, we used an optogenetic approach to demonstrate that expression of genes within region A is necessary to complete fermentations under low nitrogen availability. Overall, our results support the idea that region A is a genetic signature for wine yeast strains adapted to low nitrogen conditions.

## INTRODUCTION

Nitrogen availability in the grape must is one of the main determinants of yeast biomass generation and fermentation rate in the winemaking process (1). In the grape must, the Yeast Assimilable Nitrogen (YAN) is compound by ammonium and amino acids (2), being necessary an approximative concentration of 140 mg N/L to complete fermentation (3, 4). Below this YAN limit, the reduction in the fermentation rate generates sluggish or stuck fermentations (5, 6), leading to incomplete fermentations with high levels of residual sugar (glucose and fructose). This problem is solved by the winemaking industry supplementing the wine must with ammonium salts, which generates a negative effect on the volatile compound production (2). Importantly, the YAN necessary to complete fermentation is also dependent on the nitrogen demand of the wine yeast strain used in the fermentation process, fluctuating from 120 to 300 mg N/L (2, 7). Therefore, nitrogen availability in the grape must and nitrogen consumption by the wine yeast strains are important parameters in the winemaking process.

The importance of wine yeast strains of *Saccharomyces cerevisiae* during the fermentation process has led to the genome sequencing of hundreds of strains (8–11). For instance, the 1002 yeast genomes project sequenced 362 yeast strains that belongs to the Wine European cluster (10). Interestingly, the genome sequencing of wine yeast strains has revealed a variety of mechanisms underlying niche-specific adaptations, including structural variations, introgressions, Horizontal Gene Transfer (HGT), and hybridization (11, 12). Among these genetic characteristics, HGT are commonly absent in the yeast reference genome (laboratory strain S288C) and have typically been acquired from a distant species (10, 11). In general, HGT occurs between yeast species that share an ecological niche, and while the mechanisms are largely unclear (13), they likely involve Ty1 elements (14) and the generation of new telomeres at the chromosome ends (15).

Among wine yeasts, analysis of the genome sequence of the EC1118 commercial strain revealed three regions that were absent in the reference genome, termed regions A, B, and C (16). These regions include 34 genes involved in key functions for wine fermentation such as nitrogen and carbon source utilization (16), and many of them, particularly those in region C, have been characterized (17–20). For instance, the study of genes in region C have demonstrated the importance of the *FSY1* gene, which has been functionally characterized as a fructose transporter (17). Region C also includes open reading frames (ORFs) related to oligopeptide transport (*FOT* genes), which confer an important adaptative advantage, expanding the sources of nitrogen that can be utilized by wine yeasts during grape must fermentation (18, 19). Furthermore, region C contains the *XDH1* gene, which encodes for a xylitol dehydrogenase necessary for yeast growth on xylose (20). While many genes in region C have been characterized, genes in region A have received comparatively less attention, even though this region is also implicated in fermentative phenotypes (11, 16, 21). Thus, functional characterization of such genes may provide relevant information into yeast adaptation to the fermentative process.

Region A encodes 12 genes related to carbon and nitrogen metabolism and is mainly present in wine yeast strains (11, 16). Indeed, the ‘ALL’ (standardized name in the 1002 yeast genome project (10)) wine yeast strain carries a single copy of region A and lacks other horizontally acquired regions (21). Furthermore, the genes inside region A are transcriptionally active in this strain under fermentative conditions, and deletion of ORF-A9, which encodes a putative thiamine (B1 vitamin) transporter, results in a reduced fermentation rate in synthetic must (21). Beyond this example, our understanding of how region A contributes to wine fermentation is limited. In this sense, the generation of a collection of yeast strains that overexpress the genes in region A can shed light on the role of these genes in fermentation.

A variety of synthetic biology tools are currently available for the precise control of gene expression in a light-dependent fashion (22). Among these, natural photosensitive proteins from a variety of organisms have been used to develop optogenetic switches, permitting the control gene expression in multiple organisms, including yeast (23–25). The FUNgal Light Oxygen Voltage (FUN-LOV) optogenetic switch allows for high levels of gene expression upon blue-light stimulation and low background signal in the dark (26). This switch is based on the light-dependent interaction of LOV domains from the White Collar 1 (WC-1) and Vivid (VVD) proteins from the filamentous fungus *Neurospora crassa*, activating target genes placed under the control of the *GAL1* promoter (26). Recently, the original FUN-LOV system was redesigned by placing its components in a single plasmid (FUN-LOV^SP^ variant), using different strong promoters, and by including an antibiotic resistance as selectable marker (FUN-LOV^SP-Hph^ variant) (27). This variant was recently integrated into the *HO* locus of 59A-EC1118, a wine yeast strain, where we have demonstrated its functionality by reporting high levels of light-activated expression of a luciferase reporter gene (27). Thus, the FUN-LOV^SP-Hph^ variant can be used for the functional characterization of horizontally acquired genes in wine yeast strains to study their role in the fermentation process.

In this work, we used the FUN-LOV^SP-Hph^ variant to control gene expression of 12 genes in region A of the ‘ALL’ wine yeast strain. Initially, we determine the location of region A in the genome of the ‘ALL’ strain using long-read sequencing. Then, we generated a collection of 12 strains exhibiting light-controlled expression of each individual gene within region A. These strains were evaluated for growth parameters and fermentation kinetics in constant darkness (repression) and constant blue-light (overexpression). Our results show that expression of genes within region A is necessary to complete fermentation under low nitrogen conditions. Overall, our optogenetics-based approach highlights the importance of region A in wine fermentation and shed light on yeast adaptation to nitrogen-limited environments.

## RESULTS

### Structure and optogenetic control of region A in a wine yeast strain

We selected the ‘ALL’ wine yeast strain since its genome was previously sequenced using short reads as part of the 1002 yeast genomes projects (10), showing that region A is present in a single copy, although its physical location remains uncertain. To identify the genomic location of region A, we sequenced the genome of the ‘ALL’ strain using long-read DNA sequencing (see Materials and Methods). We obtained a close telomere-to-telomere genome assembly, confirming that region A is in a single copy, and tracked its location to the subtelomeric regions of either chromosome (Chr) VI or Chr VIII (Fig. 1). As previously reported, we found region A to contain 12 predicted genes, which are located next to the *IMD2* (YHR216W) and *PHO12* (YHR215W) genes (Fig. 1). In the S228C strain, *IMD2* and *PHO12* are the last annotated genes in the right subtelomeric region of Chr VIII. Interestingly, in the ‘ALL’ strain, *IMD2* and *PHO12* are also detected in Chr VI left subtelomeric region, suggesting a possible translocation between Chr VI and Chr VIII.

**FIG 1.**
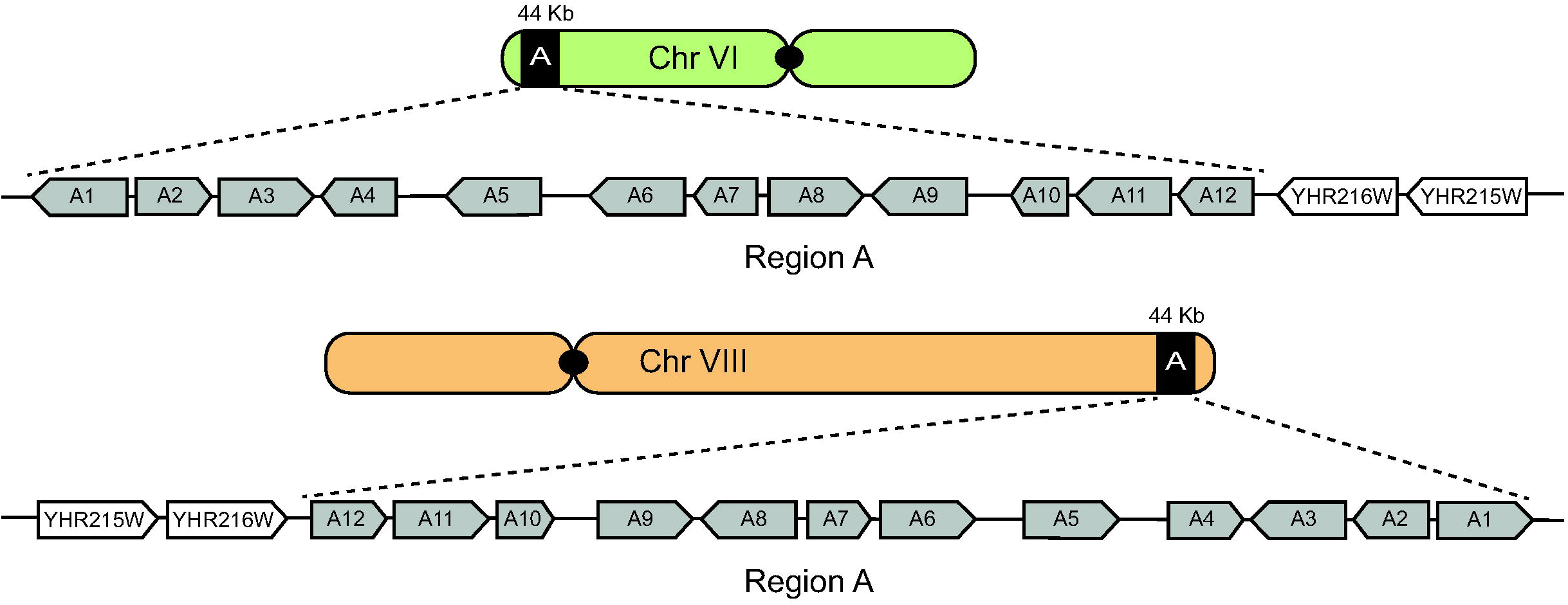
Region A location in the ‘ALL’ wine yeast strain genome. The region A is located in the subtelomeric region of chromosome (Chr) VI or VIII, next to YHR216W and YHR215W genes. Region A contains 12 predicted ORFs as previously described by (16).

After determining the genomic location of region A, we evaluate whether it was feasible to implement an optogenetic system in the ‘ALL’ strain. The FUN-LOV system is based on the light-dependent reconstitution of the Gal4 transcription factor, allowing gene expression of target genes placed under *GAL1* promoter elements (Fig. 2A). The FUN-LOV^SP-Hph^ variant was integrated into the *HO* locus and the correct functioning of this optogenetic switch was assayed using luciferase (*Luc*) and super- folder Green Fluorescent Protein (*sfGFP)* as reporter genes (Fig. 2B). These reporter genes were individually integrated into the endogenous *GAL3* locus under either the control of the *GAL1* promoter (*P_GAL1_*) or the *5XGAL1* synthetic promoter (*P_5XGAL1_*) (Fig. 2B). We decided to perform these experiments with integrated reporters (rather than episomal ones) to better emulate regulation of horizontally acquired genes in the “ALL” wine yeast strain.

**FIG 2.**
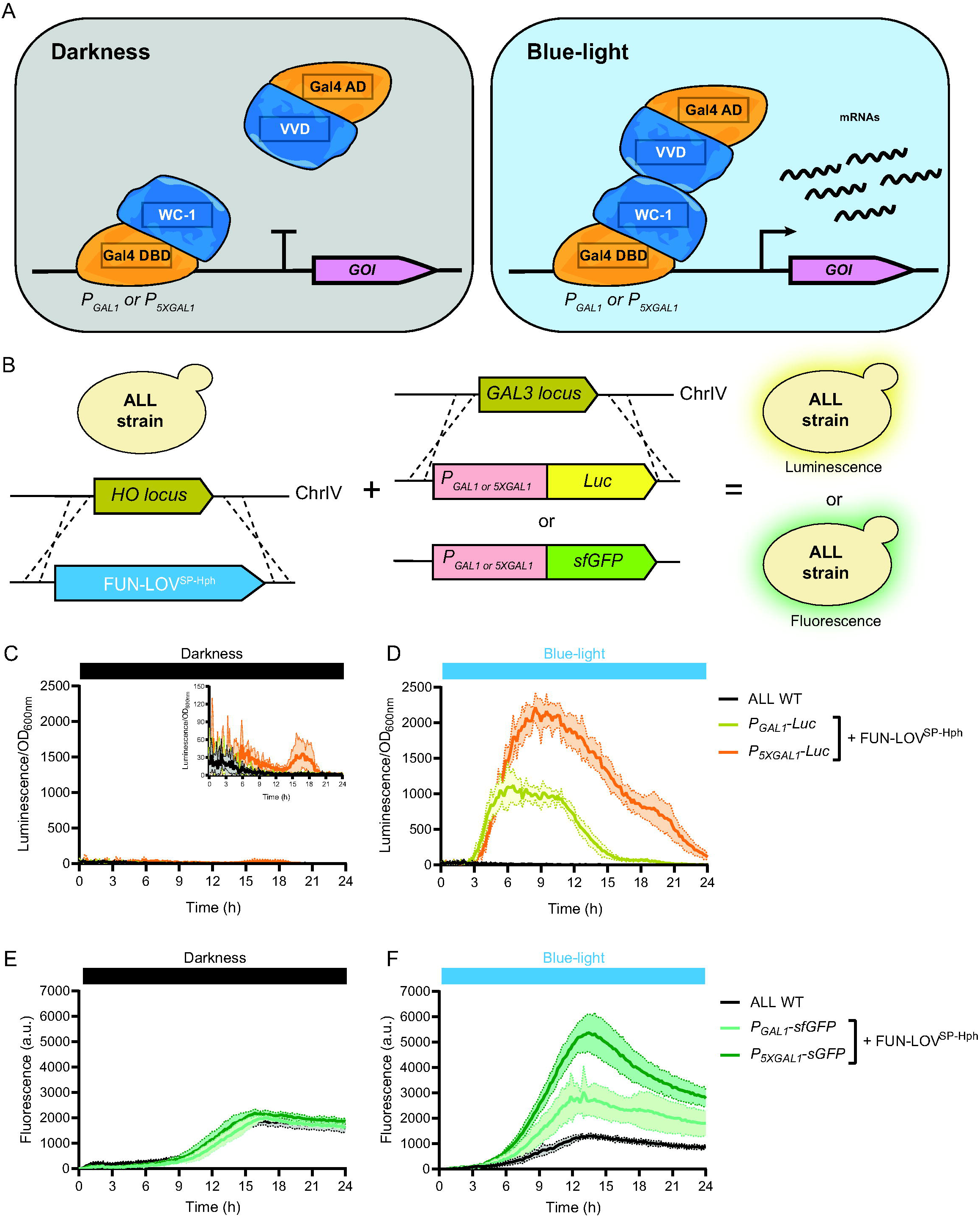
The FUN-LOV^SP-Hph^ system is functional in the ‘ALL’ wine yeast strain. **(A)** Architecture of the FUN-LOV optogenetic switch. The Gal4 DNA-Binding Domain (DBD) is linked to the LOV domain of WC-1, and the Gal4 Activation Domain (AD) is linked to the LOV domain of VVD. Upon blue-light stimulation, the interaction between the LOV domains reconstructs the Gal4 transcriptional factor, resulting in the expression of the target Gene of Interest (GOI) placed under control of *GAL1* regulatory elements. **(B)** The ‘ALL’ wine yeast strain carrying the FUN-LOV^SP-Hph^ in the *HO* locus. Further, the luciferase (*Luc)* or super-folder Green Fluorescent Protein (*sfGFP*) reporter genes were integrated into the *GAL3* locus of the same strain and are controlled by either *GAL1* (*P_GAL1_*) or *5XGAL1* (*P_5XGAL1_*) promoters. **(C and D)** *Luc* expression controlled by the FUN-LOV^SP-Hph^ variant in the ‘ALL’ strain. *Luc* was measured as luminescence and normalized by the Optical Density at 600 nm (OD_600nm_) of the corresponding yeast culture in constant darkness (left) and constant blue-light (right) conditions. **(E and F)** *sfGFP* expression controlled by the FUN-LOV^SP-Hph^ variant. sfGFP was measured as fluorescence of the yeast cultures in constant darkness (left) and constant blue-light (right) conditions. For C-F, the average of six biological replicates is shown, with the standard deviation represented as color shaded regions.

We first measured optical density at 600 nm (OD_600nm_) and luminescence (*Luc* expression) of the yeast cultures under constant darkness (DD) and constant blue-light (BL). As expected, we observed low background levels of *Luc* expression in DD (Fig. 2C) and high levels of *Luc* expression in BL (Fig. 2D). Furthermore, *Luc* expression levels were higher when the FUN-LOV^SP-Hph^ optogenetic switch was coupled to the *P_5XGAL1_* compared to the *P_GAL1_* promoter (Fig. 2D). However, background *Luc* expression in DD was higher for the *P_5XGAL1_*(Fig. 2C). Analogously, *sfGFP* expression was assayed under DD and BL conditions for the ‘ALL’ strain carrying FUN-LOV^SP-Hph^ (Fig. 2E, F). Similar to the situation with luciferase, we observed higher fluorescence levels when the FUN-LOV^SP-Hph^ targeted the *P_5XGAL1_* promoter in BL (Fig. 2F). However, no differences were observed for both promoters in DD (Fig. 2E). In parallel, we additionally performed fluorescence microscopy for the generated strains and, compared the fluorescence of the microcultures at the end of the incubation period, in both DD and BL conditions (Fig. S1-S2). Fluorescence microscopy confirmed the functionality of the FUN-LOV^SP-Hph^ optogenetic switch in the ‘ALL’ strain (Fig. S1). Furthermore, quantification of the fluorescence signal also showed that the background of *sfGFP* expression in DD is higher for the *P_5XGAL1_* compared to *P_GAL1_* (Fig. S2). Overall, the results show that the FUN-LOV^SP-Hph^ variant allows the light-controlled gene expression of a reporter gene (*Luc* or *sfGFP*) under control of *GAL1* regulatory elements in the ‘ALL’ wine yeast strain.

### Optogenetic control of genes within the horizontally acquired region A reveal their importance for yeast growth in nitrogen-limited synthetic must

The background gene expression levels observed in DD for the reporter genes controlled by the *P_5XGAL1_* through the FUN-LOV^SP-Hph^ variant suggest that this promoter may not be suitable for the optogenetic control of target genes in this strain, including those in the A region, since the DD condition may not be representing repression of these genes. To evaluate this directly, we replaced the endogenous promoter regions of ORF-A6 and ORF-A8 by the *P_GAL1_* and *P_5XGAL1_* promoters in the ‘ALL’ wine yeast strain carrying the FUN-LOV^SP-Hph^ variant (Fig. 3A). We selected these genes because they have been described as the most transcriptionally and translationally active under low nitrogen conditions (21). We then quantified the expression of ORF-A6 and ORF-A8 by RT-qPCR, and observed higher mRNA levels for these genes under BL for both promoters (*P_GAL1_* and *P_5XGAL1_*) (Fig. 3B, C). As expected, the use of *P_5XGAL1_*resulted in higher levels of gene expression compared to *P_GAL1_*under BL condition for both ORF-A6 and ORF-A8 (Fig. 2B, C). However, the increase in gene expression obtained using the *P_5XGAL1_*in BL also resulted in higher background levels of gene expression in DD (Fig. 3B, C). This background expression in DD for the *P_5XGAL1_* is even higher than the wild-type gene expression levels of ORF-A6 and ORF-A8 under their endogenous promoters (Fig. 3B, C). Conversely, when ORF-A6 and ORF-A8 are controlled by *P_GAL1_*, we observed upregulation of these genes in response to BL and, importantly, low mRNA levels in DD that are comparable to the wild-type levels of gene expression (Fig. 3B, C). Therefore, we used *P_GAL1_* to control the expression of genes inside region A and assess their contribution to different phenotypes.

**FIG 3.**
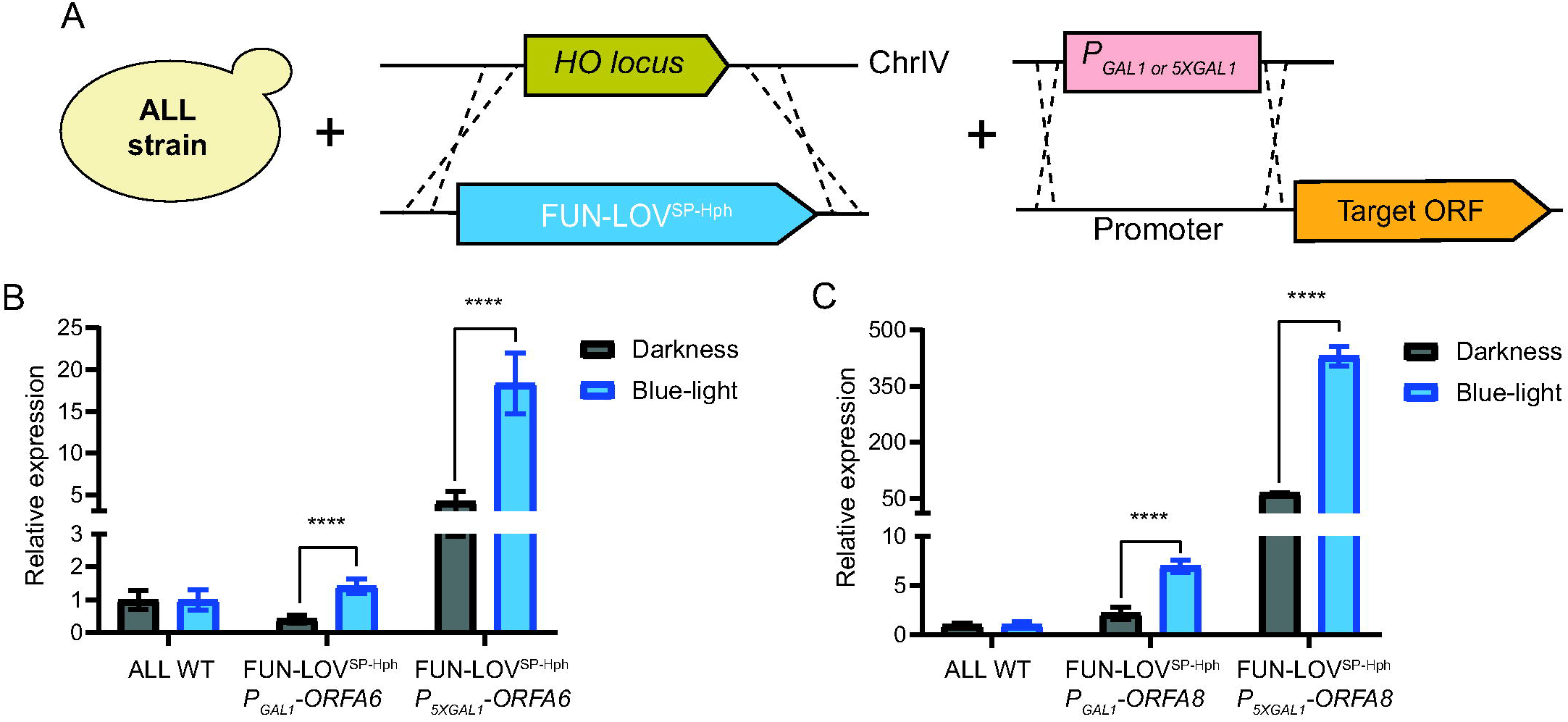
Optogenetic control of genes within region A via FUN-LOV^SP-Hph^. **(A)** The ‘ALL’ wine yeast strain carrying the FUN-LOV^SP-Hph^ in the *HO* locus and *GAL1* (*P_GAL1_*) or *5XGAL1* (*P_5XGAL1_*) promoters controlling the target ORF inside region A. (**B and C**) Light-controlled expression of ORF-A6 (**B**) and ORF-A8 (**C**) measured by RT-qPCR under constant darkness and constant blue-light. The average of six biological replicates is shown, with the standard deviation represented as error bars. The asterisks represent a statistically significant difference between constant darkness and constant blue-light conditions (*t*-test, **** = p < 0.0001).

We generated a collection of 12 strains with the *P_GAL1_* controlling the expression of each of the genes within region A, using the ‘ALL’ strain carrying the FUN-LOV^SP-Hph^ variant (Fig. 3A). To study the contribution of these genes for growth under fermentation-relevant conditions, we evaluated the growth of these strains under DD and BL conditions using Synthetic Must (SM) with different nitrogen availability: 60 mg/L (SM60), 140 mg/L (SM140), and 300 mg/L (SM300) (Fig. 4). At SM300 (high nitrogen concentration) and SM140 (control), the collection exhibited similar kinetic profiles in both DD and BL conditions (Fig. 4A, B, C, D), with exception of the ORF-A8 at SM300 (raw data in Fig. S3-S6). Interestingly, under nitrogen limitation (SM60), a large difference was observed in the growth kinetics in DD compared to the BL condition (Fig. 4E, F; raw data in Fig. S7-S8). These results suggest that expression (in BL) of region A genes may be beneficial for proper growth under nitrogen-limited synthetic must (Fig. 4E, F). To quantitatively compare the observed growth differences in DD respect to BL for the tested strains, we calculated the Area Under the Curve (AUC) for each growth profile (28), normalizing the data by the AUC of the wild type “ALL” strain in BL and DD (see Materials and Methods). These results confirmed the differences in DD respect to BL for SM60 growth kinetics (Fig. 4G). In conclusion, the use of optogenetic-controlled gene expression within region A demonstrates its contribution in improving the kinetics of growth under low nitrogen conditions.

**FIG 4.**
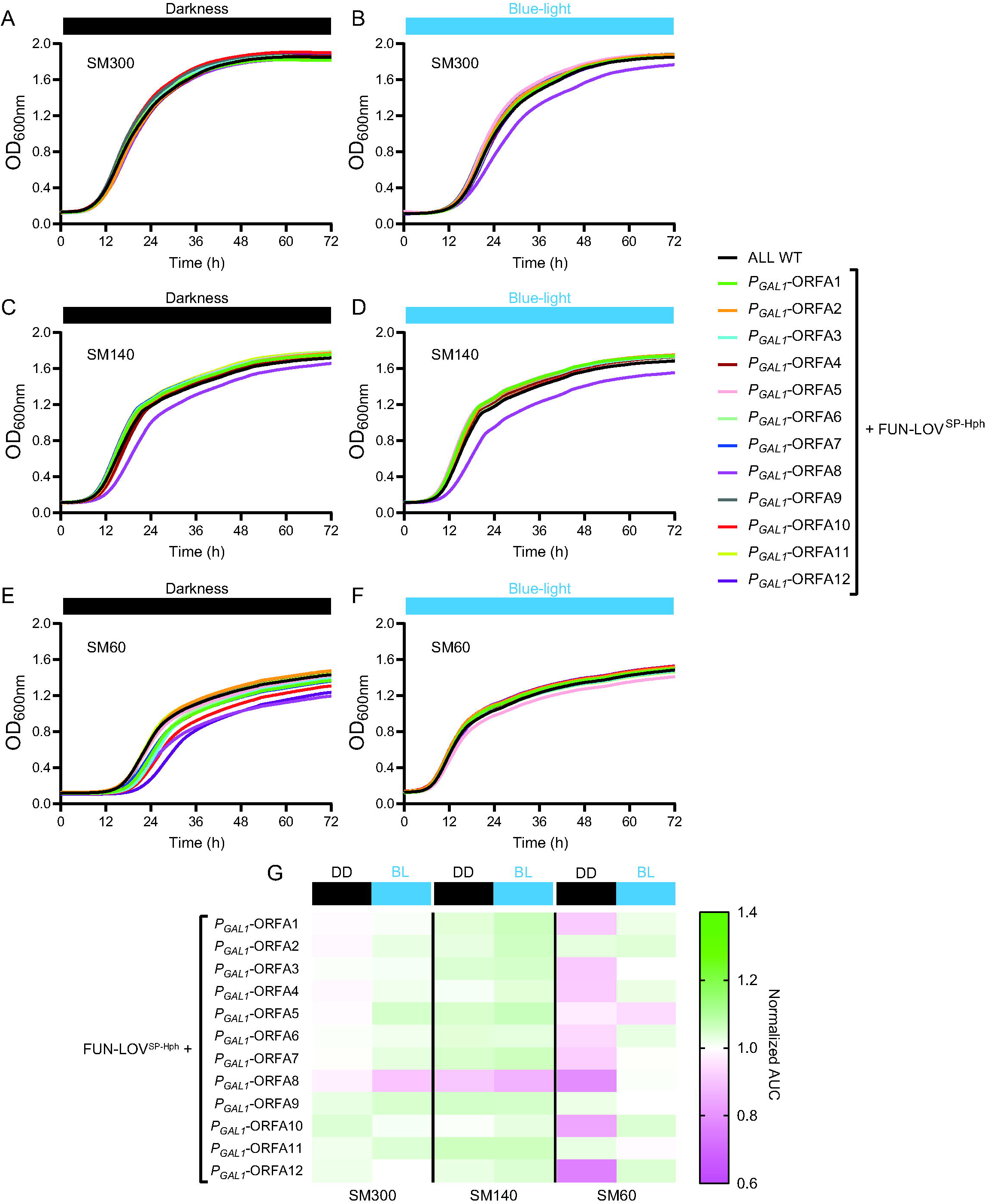
Modulation of gene expression for genes within region A affects growth kinetics in synthetic must with different nitrogen availability. The growth curve of the ‘ALL’ wild-type strain was compared with strains carrying the FUN-LOV^SP-Hph^ variant and the *GAL1* promoter (*P_GAL1_*) controlling the expression of different genes within region A. (**A and B**) Growth curves in SM300 under constant darkness and constant blue-light conditions, respectively. **(C and D)** Growth curves in SM140 under constant darkness and constant blue-light conditions, respectively. **(E and F)** Growth curves in SM60 under constant darkness and constant blue-light conditions, respectively. **(G)** Area Under the Curve (AUC) extracted from the growth curves shown in panels **A** to **F**. AUC was normalized by dividing the phenotype (AUC) of each strain by the phenotype (AUC) of the ‘ALL‘ wild-type strain in constant darkness (DD) and constant blue-light (BL) conditions, and is represented as a heatmap. The average of six biological replicates is shown.

### Activation of genes within region A prevent stuck fermentations in nitrogen-limited synthetic must

Our data suggested that genes in region A are relevant for growth of the ‘ALL’ strain under nitrogen-limited conditions, a situation that is common during fermentation. To directly evaluate the importance of these genes during fermentation, we performed laboratory-scale fermentations under DD and BL conditions. We assayed all 12 strains, each with *P_GAL1_* controlling the expression of one of the genes within region A, in SM60, SM140, and SM300, measuring CO_2_ loss at multiple times during fermentation (Fig. 5).

**FIG 5.**
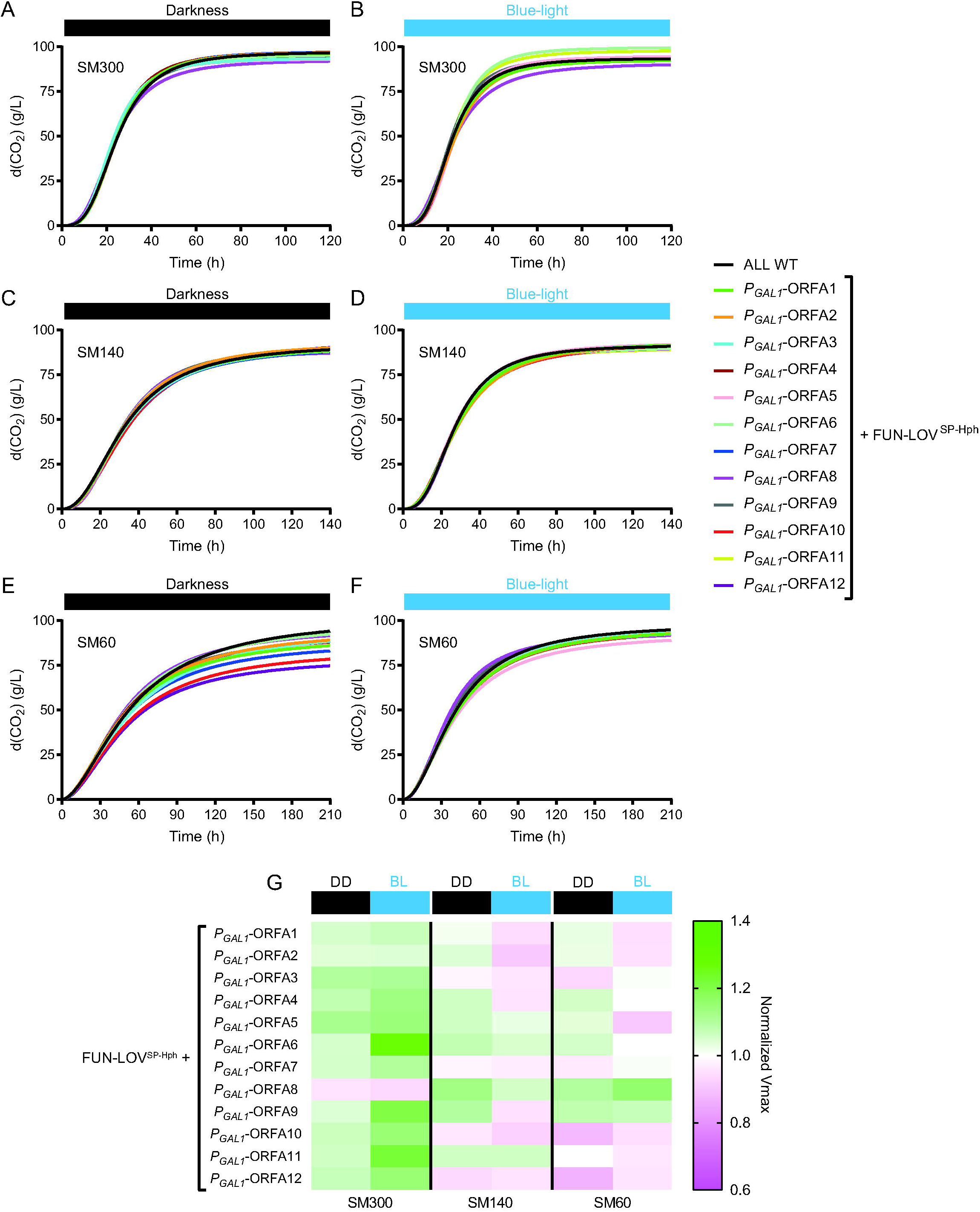
Modulation of gene expression for different genes within region A affects fermentation in low- nitrogen synthetic must. The CO_2_ release of the ‘ALL’ wild-type strain was compared with that of strains carrying the FUN-LOV^SP-Hph^ variant and the *GAL1* promoter (*P_GAL1_*) controlling the expression of different genes within region A. (**A and B**) Fermentation kinetics in SM300 under constant darkness and constant blue-light conditions, respectively. **(C and D)**, Fermentation kinetics in SM140 under constant darkness and constant blue-light conditions, respectively. **(E and F)**. Fermentation kinetics in SM60 under constant darkness and constant blue-light conditions, respectively. In the fermentation kinetic plots, the CO_2_ release curves were fitted to a sigmoid non- linear regression. **(G)** Maximal CO_2_ production rate (V_max_) extracted from the fermentation kinetic profiles shown in panels A to F. V_max_ was normalized by dividing the phenotype (V_max_) of each strain by the phenotype (V_max_) of the ‘ALL’ wild-type strain in constant darkness (DD) and constant blue- light (BL) conditions, and is represented as a heatmap. The average of three biological replicates is shown.

Fermentation kinetics for all strains in SM300 and SM140 showed no significant differences between DD and BL conditions (Fig. 5A, B, C, D; raw data in Fig. S9-S12). In contrast, analysis of the fermentation kinetics under SM60 revealed a decrease in CO_2_ loss under DD for several strains, a phenotype that was restored under the BL condition (Fig. 5E, F; raw data in Fig. S13-S14), which was consistent with the growth curve results (Fig. 4). We further compared the DD and BL fermentation phenotypes by extracting the maximal fermentation rate (V_max_) from the CO_2_ loss curves (Fig. S15) and normalizing the data with the V_max_ of the ‘ALL’ wild type strain under DD and BL (see Materials and Methods). The results confirmed that upregulation (in BL) of different ORFs within region A change the V_max_ (compared to DD) in SM60 (Fig. 5G). For instance, in SM60, deactivation of ORF-A10 and ORF-A12 in DD results in a decreased V_max_, but their overexpression in BL increases the V_max_ respect to the wild-type strain (Fig. 5G). On the contrary, also in SM60, ORF-A5 overexpression in BL decreases the V_max_ and its repression in DD increases the V_max_ compared to the wild-type “ALL” strain (Fig. 5G). Therefore, expression of multiple genes within region A contribute to maintaining the fermentation rate under low nitrogen conditions in a wine yeast strain.

The phenotypic differences observed in the fermentation rate for DD and BL conditions in the generated strains, suggest that the main products of the fermentation and residual sugar could be affected by repression/overexpression of the genes within region A. Thus, we used high- performance liquid chromatography (HPLC) to measure the main fermentation products (glycerol, ethanol, and acetic acid) and sugar consumption (glucose and fructose) at the end of the fermentation process.

For the metabolites, and to compare the DD and BL conditions, the concentration of each of the metabolites was normalized by that observed in the wild type ‘ALL’ strain in each condition. The results show that the metabolite profile changes in DD compared to BL in all the fermentation conditions tested (SM60, SM140, and SM300), with most differences observed in SM60 and SM140 (Fig. 6A).

**FIG 6.**
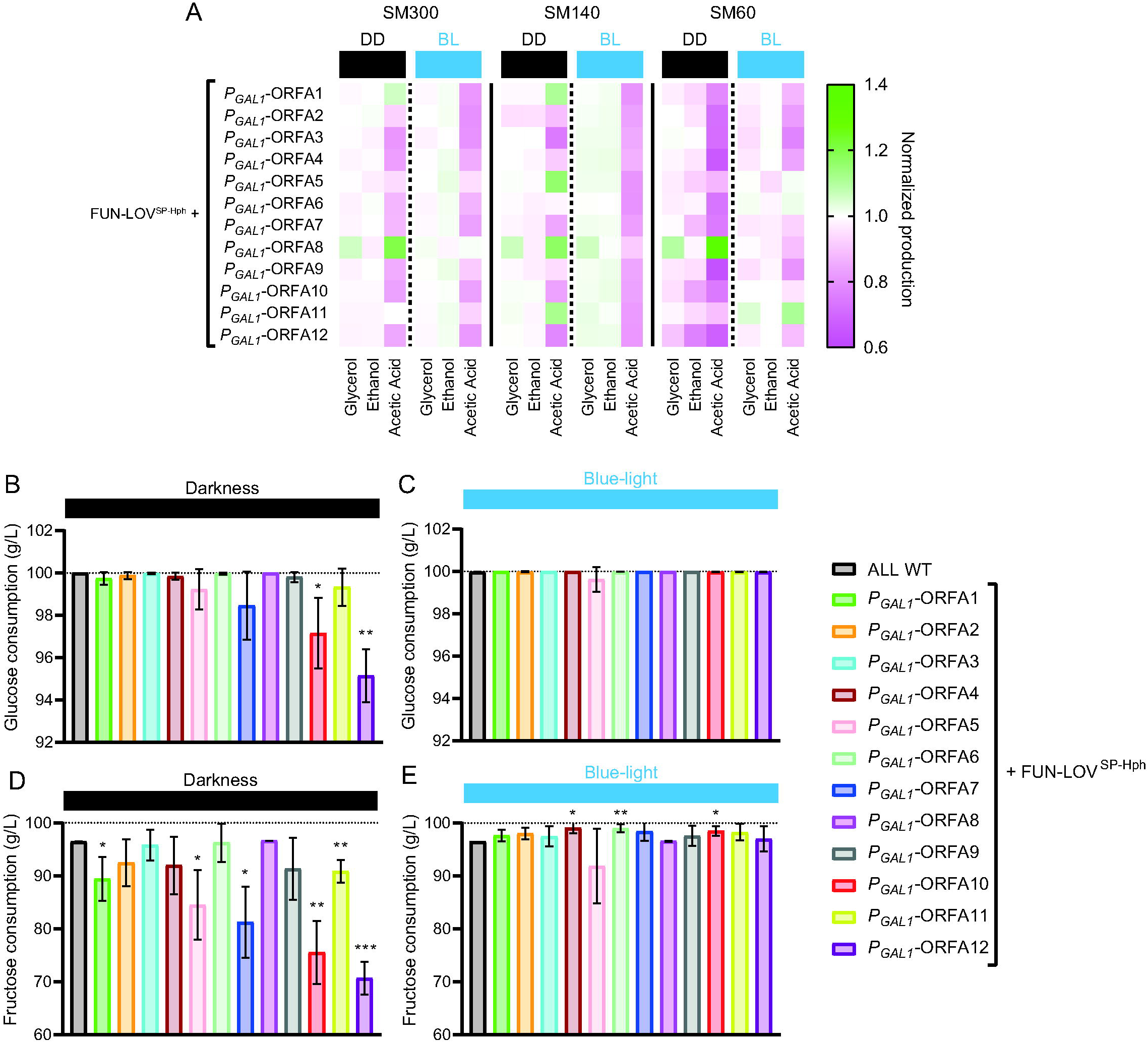
The expression of genes within region A prevents stuck fermentations in low-nitrogen synthetic must. **(A)** Fermentation by-products (glycerol, ethanol, and acetic acid) were measured at the end of fermentations in SM300, SM140, and SM60. The metabolite production (concentration) was normalized by dividing the values from each strain by the phenotype (metabolite production) of the ‘ALL’ wild-type strain and are represented as a heatmap in constant darkness (DD) and constant blue-light (BL) conditions. **(B – E)** Sugar consumption for fermentations in SM60. (**B and C**) Glucose consumption under constant darkness and constant blue-light, respectively. (**D and E**) Fructose consumption under constant darkness and constant blue-light, respectively. Dashed lines represent the initial values of glucose and fructose. The average sugar consumption of three biological replicates with the standard deviation represented as error bars is shown. The asterisks represent a statistically significant difference respect to the ‘ALL’ wild-type strain (*t*-test, * = p < 0.05; ** = p < 0.01).

We then measured (by HPLC) the sugar consumption of the strains at the end of the fermentations in SM60, SM140, and SM300, under DD and BL conditions. Interestingly, several strains were unable to consume the available glucose and fructose in SM60 under DD compared to the wild-type strain (Fig. 6B, C; Fig. S16), suggesting incomplete fermentations. Conversely, fermentations in SM60 under BL condition showed full glucose consumption and an increase in fructose consumption for various strains (Fig. 6D, E). For instance, overexpression in BL of ORFs A7, A10, and A12 increased glucose and fructose consumption compared to DD (compare Fig. 6B to 6D; and Fig. 6C to 6E), suggesting that those genes are important for fermentation under low nitrogen conditions. In conclusion, our results in SM60 demonstrate that upregulation of genes within region A can prevent stuck fermentations under low nitrogen conditions, suggesting that they may play a role in this process.

## DISCUSSION

One of the most common problems in winemaking is low nitrogen levels in grape must (2, 6). Such low levels are typically unable to fulfill the nitrogen requirements of the wine yeast strains, leading to incomplete fermentations with high levels of residual sugar (2, 6). This problem is usually solved by nitrogen addition into the grape must, which can affect the wine aromatic profile and potentially result in the production of biogenic amines (29, 30). In this sense, our results show that expression of genes within region A (e.g. ORF-A7, ORF-A10, and ORF-A12) promotes fermentation completion in low nitrogen synthetic must (Fig. 5), which is associated with a reduced amount of residual sugar (lower than 5 g/L) at the end of the fermentation process (Fig. 6). Conversely, repression of these genes, was associated with reduced sugar consumption and lower amounts of the main fermentation products (ethanol, glycerol, and acetate) in nitrogen-limited synthetic must (Fig. 6), which is consistent with an incomplete fermentation. Further, the expression of genes from this region may be explored as potential genetic markers for selection of wine yeast strains with fermentation proficiency under nitrogen limitation.

The budding yeast *S. cerevisiae* has become a powerful biological platform for optogenetics, and there are several tools available that allow researchers to use light to control different phenotypes (23). Among the available optogenetic tools, the FUN-LOV system enables light-controlled expression of the targeted genes (26). This system has been used to regulate different yeast phenotypes, including flocculation, heterologous protein production, and the mating type signaling pathway (26, 31). In this work, we assess the contribution of horizontally acquired genes in fermentation. To do this, we used a previously described FUN-LOV variant (FUN-LOV^SP-Hph^) to control the expression of the horizontally acquired genes of region A in a yeast wine strain and demonstrated the importance of this region in wine fermentation under low nitrogen availability.

We first sequenced the genome of the ‘ALL’ wine yeast strain using a long-reads approach, identifying that region A is in a single copy with two possible subtelomeric locations (Fig. 1). Interestingly, the potential location of region A at the beginning of ChrVI resembles the previously described location of this region in the EC1118 wine yeast strain (16). We then integrated the FUN- LOV^SP-Hph^ variant into the genome of ‘ALL’ strain and confirmed its functionality using *Luc* and *sfGFP* as reporter genes (Fig. 2). The gene expression levels achieved by the FUN-LOV^SP-Hph^ variant were consistent with those previously reported in the 59A-EC1118 wine yeast strain (27). We then used the FUN-LOV^SP-Hph^ variant for the light-dependent control of gene expression of 12 genes within region A, and confirmed the expected upregulation in response to light for ORF-A6 and ORF-A8 by RT-qPCR (Fig. 3). The advantage of this optogenetic strategy is that it allows activation (by light) or deactivation (by dark) of gene expression in the same yeast strain, allowing the comparison of phenotypes across different fermentation conditions. Through such an optogenetic tool, we attempted to functionally characterize the contribution of horizontally acquired genes within region A, in yeast fermentation.

Region A has been reported to be mainly present in wine yeast strains, and it has been horizontally acquired from *Torulaspora sp.* (10, 11, 21). A few genes within this region A have been reported to play important roles in yeast adaptation to fermentative environments, with putative functions related to carbon and nitrogen metabolism (16, 21). To further study the role of these genes in wine fermentation, we performed growth curves (Fig. 4) and laboratory-scale fermentations (Fig. 5) for yeast strains carrying each gene of region A individually controlled by the FUN-LOV^SP-Hph^ variant, thus allowing for the modulation of their expression by blue light. Our results show that the expression of different genes within region A contributes to yeast growth and fermentation under nitrogen-limited synthetic must. For instance, modulating the expression of ORFs A10 and A12 resulted in major impacts on growth kinetics and fermentation rate in SM60 (Fig. 4G, 5G). These genes encode a putative methyltransferase involved in ubiquinone biosynthesis (*COQ5*), and a possible 2-nitropropane dioxygenase or nitronate monooxygenase (*YJR149W*), respectively (16, 21).

Ubiquinone, also known as coenzyme Q (CoQ), is an important component of the electron transport chain and regenerates cellular antioxidants. In *S. cerevisiae,* expression of the *COQ5* gene is regulated by carbon sources (32). Furthermore, expression of CoQ biosynthesis genes is also modulated downstream by other regulatory mechanisms such as mitochondrial protein import, assembly of the CoQ protein complex, and phosphorylation cycles that regulates the last steps of CoQ biosynthesis (33). Although glucose fermentation does not require a functional mitochondrial electron transport (34), additional non-mitochondrial functions for CoQ have been identified, such as regeneration of ascorbate free radicals at the plasma membrane (35). Interestingly, yeast strains mutant for the *COQ5* gene show a decreased rate of nitrogen source utilization (36) and a reduced rate of vegetative growth (37). These previously described observations are consistent with our results for ORF-A10, which show lower growth kinetics and fermentation rate in dark (deactivation) compared to the blue-light condition (overexpression) in low nitrogen conditions (Fig. 4, 5). On the other hand, ORF-A8 showed an abnormal behavior compared to the wild-type strain for growth kinetics assays and metabolite production at the end of fermentation (Fig. 4, 6). This gene encodes a possible member of the multi-drug and toxin extrusion family (*ERC1*) (16, 21). *ERC1* overexpression confers ethionine resistance, accumulation of S-adenosylmethionine (SAM), and decreased rate of vegetative growth in yeast (37, 38), while *ERC1* mutation increases competitive fitness in minimal medium (39). These reports are consistent with our results for growth curves in SM300, where constant darkness (deactivation) increased growth parameters compared to blue light (overexpression) (Fig. 4). Overall, although the molecular mechanism concerning how genes from region A are integrated into the yeast metabolism is still unclear, our results demonstrate their importance for fermentation in low nitrogen conditions.

In conclusion, we used an optogenetic tool in a wine yeast strain, which enabled the uncovering of the contribution of horizontally acquired genes in yeast fermentation. Indeed, we show that expression of genes within region A contributes both to maintain proper fermentation kinetics and prevent stuck fermentations under nitrogen-limited conditions. Overall, our results open important questions concerning how genes from region A interplay with genes of the core yeast genome, permitting to overcome nitrogen limitations and to complete wine fermentation. Finally, our results highlight the impact of horizontally acquired genes in yeast adaptation to domesticated environments such as wine fermentation.

## Materials and Methods

### Strain generation and culture conditions

The ‘ALL’ (standardized name) wine yeast strain (isolate CYY 021-004-099; Culture Collection of Yeast) used in this work was part of the 1002 yeast genomes project (10). This strain belongs to the Wine/European cluster, and it was isolated from the Danube River in Bratislava, Slovakia (10). The ‘ALL’ strain carries a hemizygotic region A composed of 12 ORFs (21). The ‘ALL’ strain was first used for transformation and genome integration of the FUN-LOV^SP-Hph^ variant at the *HO* locus as previously described by Figueroa et al., (27). Then, the ‘ALL’ strain was transformed with the firefly luciferase (*Luc*) or super-folder Green Fluorescent Protein (sf*GFP)* reporter genes under control of either the *GAL1* promoter or the *5XGAL1* synthetic promoter (five repetitions of Gal4 Upstream Activating Sequence) (26). The reporter genes were integrated into the *GAL3* locus by PCR amplification of the construct, followed by transformation, and direct recombination with the target locus (26). In parallel, the endogenous promoter of each ORF in the region A was replaced by the inducible *GAL1* promoter or the *5XGAL1* synthetic promoter (26). Promoter swapping was performed by PCR amplification of the promoter region, transformation, and direct recombination with the endogenous promoter. All PCR amplifications for genome integration and promoter swapping were carried out with Phusion Flash High-Fidelity Master Mix (ThermoFisher Scientific, USA).

The strains were maintained on YPDA medium (2% glucose, 2% peptone, 1% yeast extract, and 2% agar) at 30 °C. Yeast transformations were carried out using the standard Lithium acetate transformation protocol (40). Transformant strains were selected on YPDA medium supplemented with 0.3 mg/mL of hygromycin or 0.4 mg/mL of G418 as required. Finally, transformant strains were confirmed by standard yeast colony PCR using GoTaq Green Master Mix (Promega, USA). The yeast strains used and generated in this work are listed in Table S1.

### Wine yeast strain genome sequencing, assembly, and annotation

The genome of the ‘ALL’ wine yeast strain was sequenced using PacBio technology through the sequencing facilities of the University of Valencia, Spain. PacBio sequencing reads were obtained using the Circular Consensus Sequencing (CCS) method. The genome of the ‘ALL’ isolate was assembled *de novo* in a highly continuous DNA sequence (16 chromosomes and 12 scaffolds) with the LRSDAY (v.1.7.2) pipeline (41, 42). We provided as input both high-coverage PacBio HiFi long reads and Illumina PE short reads (preparation steps 00) (10). PacBio HiFi long reads are available at the European Nucleotide Archive under the code PRJEB73885, and the Illumina PE short reads are available under the code ERP014555. The PacBio HiFi long reads, and the Illumina PE short reads were quality assessed with *fastqc* (v.0.11.9). The long read-based genome assembly was performed with *flye* assembler (v.2.9.1-b1780) (step 01) (43). The quality of the raw assembly was improved by running 3 successive rounds of polishing with the Illumina short reads (step 03). The polished assembly was scaffolded with the reference-based scaffolder *ragtag* (v.2.1.0) (44), using S288C whole-genome assembly as a guide (ASM205763v1) (step 04). The centromeric profiling was performed with *exonerate* (v.2.2.0) (45) and the final assembly was obtained after a manual reordering and renaming of each sequence (steps 05, 07, 08). We performed annotations of centromeres, protein-coding, and tRNA genes with *maker* (v.3.00) (46) and *EVM* (v.1.1.1) (47) (steps 08, 09). Transposable elements (TEs) and both X and Y’ chromosome elements were annotated with a custom Perl script and pre-shipped *Saccharomyces*-specific sequences with LRSDAY (steps 11, 12 and 13) (41, 42). The annotated protein-coding genes were compared with a group of references to identify gene orthologs using *proteinortho* (v.6.0.35) (48), and the SGD systematic names were assigned to the newly annotated protein-coding genes. Finally, all the annotations were integrated into a single *gff* file (steps 14, 15). The ‘ALL’ *de novo* genome assembly was quality assessed with BUSCO (v.5.6.0) (49), QUAST (v.5.2.0) (50) and MUMMer (v.4.0.0rc1) (51).

### Region A detection, location, and copy number assessment

The ‘ALL’ *de novo* genome assembly was interrogated to detect and locate the HGT region A from *Torulaspora delbrueckii*. We aligned the 12 CDS of region A to the newly assembled genome with *fasta36* (v.36.3.8h May 2020). Region A was maintained in a separated contig (contig_12) and was not assembled into a chromosome. To establish its position, we aligned the PacBio HiFi long reads back to the *de novo* genome assembly using *minimap2* (2.27-r1193) (52, 53), and the reads mapping on the flanking sides of region A were extracted. One of the flanking sides hosted a telomeric repeat and we considered it uninformative for defining the location of region A, while the protein-coding genes annotation identified the gene YHR216W on the other flank. We sketched a scheme of YHR216W and region A orientation and selected the long reads 1) spanning the lateral side of YHR216W (opposite to region A) and, 2) containing at least one gene of the HGT. In detail, we first subsetted the *.sam* file of contig_12 from position 1 to 20,000 (*samtools view*). We extracted the long reads from the *.sam* file with *samtools fasta* and created a database with *blastn* (v.2.15.0+) (54, 55). We blasted the gene YHR216W (*IMD2*) and the nearest gene of region A against the database requiring 1) 100% query covered, and 2) the two genes to be located on the same long read. We identified 2 informative long reads (m54366Ue_240214_182506/107809029/ccs and m54366Ue_240214_182506/80282722/ccs; both with MQ 60) of 14,511 bp and 13,562 bp, with YHR216W located at 10,329-11,900 bps and 9,458-11,029 bps, respectively. The ∼3-Kb flanking regions of the two long reads were investigated for the presence of additional ORFs with “ATG” as start and any stop codon of the standard code, by using the ORFfinder online NIH tool [https://www.ncbi.nlm.nih.gov/orffinder/]. We were able to identify a single ORF in both reads corresponding to YHR215W (*PHO2*), which, like YHR216W, was annotated both against chromosome VI and twice against chromosome VIII.

The number of copies of region A was estimated by short and long-read mapping against the newly assembled genome, extracting the coverage from non-overlapping 1-Kb consecutive windows. The relative coverage of region A was half the whole-genome coverage suggesting n ploidy, in contrast to the 2n ploidy of the isolate ‘ALL’. Furthermore, ‘ALL’ ploidy was previously estimated via FACS in Peter et al., (10).

### Design and generation of genetic constructs

The genetic constructs carrying the *GAL1* or *5XGAL1* promoter controlling expression of *sfGFP* and including the *KanMx* (G418) antibiotic resistance in the reverse direction (*KanMxRV*), were assembled by Yeast Recombinational Cloning (YRC) using the pRS316 plasmid as backbone (56). In the YRC assemblies, each genetic element (promoter, coding sequence, terminator, and antibiotic resistance) was PCR amplified using primers with 40 nt of overlap between fragments. All PCR reactions were carried out using Phusion Flash High-Fidelity Master Mix (ThermoFisher Scientific, USA). PCR products were then co-transformed with the linearized pRS316 plasmid into the BY4741 strain for assembly (56). The assembled plasmids were confirmed by standard yeast colony PCR and then extracted from yeast using the Zymoprep Yeast Plasmid Miniprep I (Zymo Research, USA). Selected plasmids were used for *E. coli* DH5α transformation and then confirmed by Sanger sequencing through the Eurofins sequencing service. The list of primers and plasmids used and generated in this work are listed in Tables S2 and S3, respectively.

### Luciferase expression assay

A destabilized version of the *Luc* gene was used as a reporter for light-activated gene expression (57). Real-time *Luc* assays were performed under constant darkness (DD) and constant blue-light (BL) conditions (26, 27, 58). Yeast strains harboring the constructs of interest were grown overnight in separate wells of a 96-well plate, each containing 200 µL of Synthetic Complete (SC) medium (glucose 2%, YNB w/o aa 0.68%, drop-out synthetic mix 0.2%, tryptophan 0.002%, leucine 0.01%, uracil 0.002%, and histidine 0.002%) at 30 °C. The next day, 10 µL of each of the overnight cultures was transferred to a separate well of a white 96-well plate with optical bottom (ThermoFisher Scientific, USA) containing 290 µL of SC medium and 1mM luciferin. The OD_600nm_ and luminescence of each well were acquired using a Synergy H1M microplate reader (Agilent, USA). In the DD and BL experiments, the 96-well plate was incubated outside the plate reader in a dark indoor growth system at 25 °C, but in the BL experiments, a previously described illumination system was utilized (27, 58). This illumination system provides blue light at 460 nm with 24 µmoles m^-2^ s^-1^ of light intensity (26, 27, 58). In all the experiments, the plate reader was programmed for discontinuous kinetics using the Gen5 software (Agilent, USA), measuring OD_600nm_ and luminescence of the yeast cultures every 10 min. Luciferase expression was normalized by dividing luminescence by the OD_600nm_ of the yeast cultures. All experiments were performed in six biological replicates.

### Fluorescence assays and microscopy

The *sfGFP* gene was used as a reporter for light-activated gene expression (59). Real-time fluorescence assays were performed under DD and BL conditions. Yeast strains were grown overnight in a 96-well plate format, with each well containing 200 µL of SC medium at 30 °C. The next day, 10 µL of each of the overnight cultures was transferred to a separate well of a black 96- well plate with optical bottom (ThermoFisher Scientific, USA) containing 290 µL of SC medium. The OD_600nm_ and fluorescence were acquired using a Synergy H1M microplate reader (Agilent, USA). Fluorescence measurements were performed using 485 nm for excitation and 515 nm for fluorescence acquisition. The illumination setup was identical to that described for the luciferase expression assay. All experiments were performed in six biological replicates.

Microscopy experiments and fluorescence quantification assays were performed under DD and BL conditions. Yeast strains were grown for 16 hours in a black 96-well plate, with each well containing 200 µL of SC medium at 28 °C. Fluorescence images were acquired with an Eclipse 90i epifluorescence microscope (Nikon, Japan) by using a 40X objective with a GFP filter (B-2E/C). Bright-field microscopy was used to focus on cells, and then the Nikon Digital Sight DS-5Mc was utilized for image capture under bright-field and fluorescence modes. ImageJ was used to analyze images (60). In parallel, the final OD_600nm_ and fluorescence of the yeast cultures were measured using a CLARIOstar^plus^ plate reader (BGM Labtech, Germany). Fluorescence was normalized by the OD_600nm_ of the corresponding yeast cultures. All experiments were performed in six biological replicates.

### Gene expression analysis by real-time quantitative PCR (qPCR)

Total RNA extraction and relative mRNA levels were determined as described by Bisquert et al., (61) with a few modifications. Briefly, 10-20 mL of exponentially growing yeast cells were washed with RNAse-free MiliQ water and frozen at -80 °C. Then, cell pellets were ruptured by mechanical cell lysis in a MillMix 20 homogenizer (Domel Labs, Slovenia) with 0.4 mL of LETS buffer (61) and glass beads. Supernatants were treated with phenol-chloroform (5:1) and chloroform-isoamyl alcohol (24:1). RNA was then precipitated overnight at -20 °C twice. For the first precipitation, 2.5 volumes of 96% ethanol and 0.1 volume of 5 M LiCl were added. For the second, 2.5 volumes of 96% ethanol and 0.1 volume of 3 M sodium acetate were added. Resulting RNA was resuspended in RNAse-free MiliQ water and quantified using a NanoDrop Spectrophotometer (ThermoFisher Scientific, USA). Total RNA was treated with DNAse I RNAse-free (Roche, Germany) for genomic DNA removal. cDNA was synthesized using the NZY First-Strand cDNA Synthesis kit (NZYtech, Portugal). Lastly, qPCR was performed in a Light Cycler 480 II (Roche, Germany) using Power SYBR^®^ Green PCR Master Mix (ThermoFisher Scientific, USA). Relative gene expression calculations for ORF-A6 and ORF-A8 were performed with the standard curve method described by Bisquert et al., (61) and using the *ACT1* gene as reference. The primers used in qPCR experiments are listed in the Table S2.

### Growth curves and fermentation experiments

Growth curves were performed under DD and BL conditions using Synthetic Must (SM) with different amounts of Yeast Assimilable Nitrogen (YAN) in the culture medium (62). The illumination conditions were identical as previously described above (luciferase and sfGFP experiments). The OD_600nm_ of the yeast cultures was measured using a Synergy H1M microplate reader (Agilent, USA). The yeast strains were grown overnight in a 96-well plate, with each well containing 200 µL of YNB (glucose 2%, and YNB w/o aa 0.68%) medium at 30 °C. The next day, 10 µL of each of the overnight cultures was transferred to a separate well of a 96-well plate with optical bottom (ThermoFisher, USA) containing 290 µL of SM with 60, 140, or 300 mg/N L of YAN. The Area Under the Curve (AUC) was extracted from the growth curves (28) using the GraphPad Prism 8 software (Dotmatics, USA). The AUC was normalized by dividing the phenotype of each strain by the phenotype of the wild type ‘ALL’ strain and was represented as a heatmap using GraphPad Prism 8 software (Dotmatics, USA). All growth curve experiments were performed in six biological replicates.

Laboratory-scale fermentations were assayed in 5 mL of SM with 60, 140, or 300 mg/N L of YAN, using an incubator to control the temperature at 28 °C and a magnetic stir plate at 150 rpm to homogenize the cell culture. Fermentations were inoculated with 2 x 10^6^ cells/mL, and progression was monitored by CO_2_ release, measured as weight loss during the time course of the experiments and until measurements were stable (21, 62). The CO_2_ loss curves were fitted to a sigmoid non- linear regression and the first derivative was calculated to obtain the maximal CO_2_ production rate (V_max_) of each strain (63). V_max_ values were normalized by dividing the value obtained for each strain by that of the wild type ‘ALL’ strain and were represented as a heatmap using GraphPad Prism 8 software (Dotmatics, USA). Illumination was performed using a LumiGrow Pro 650TM LED array (LumiGrow, USA), emitting blue light at 450 nm with 40 µmoles m^-2^ s^-1^ of light intensity (64). All fermentation experiments were performed in three biological replicates.

### HPLC analysis

Sugars (glucose and fructose) and fermentative by-products (glycerol, ethanol, and acetic acid) were determined by HPLC as described by Pérez et al., (65). Briefly, supernatants from fermentations were filtered using a 0.22-µm filter and diluted according to their estimated residual sugar amount. Then, samples were analyzed by HPLC using refraction index and UV/VIS (210 nm) detectors equipped with a HyperREZTM XP Carbohydrate H+ 8 mm column and HyperREZTM XP Carbohydrate Guard (ThermoFisher Scientific, USA). The analysis conditions were: 0.6 mL/min flux of H_2_SO_4_ 1.5 mM, 35 bars of pressure, and pre-heating at 50 °C. The concentration (g/L) of each compound was determined with calibration curves using the corresponding standard compound (Merck, Germany).

## DATA AVAILABILITY

The original contributions presented in the study are included in the article/Supplementary Material, further inquiries can be directed to the corresponding author.

## ACKNOWLEDGMENTS

We thank M. Teresa Lafuente Rodríguez for give us access to the LumiGrow Pro 650TM LED array equipment. We thank Alejandro Montenegro-Montero for language and editorial support.

This research was funded by ANID-Millennium Science Initiative Program-ICN17_022 and ANID- FONDECYT grant number 1210955 to FS; ANID-FONDECYT grant number 11220533 and ANID- FONDEF IDeA I+D grant number ID24I10027 to EKP; ANID-FONDECYT grant number 1201104 and ANID-FONDEF IDeA I+D grant number ID21I10198 to CM; and by the ANID-PhD scholarships 21200745 to DF. AQ and JMG thanks to the Spanish Government, ref. MCIN/AEI/10.13039/501100011033, as a IATA (CSIC) ‘Severo Ochoa’ Center of Excellence (CEX2021-001189-S) and AQ to MCIU/AEI/FEDER grant references PID2021-126380OB-C31.

DF and DR generated the plasmids, developed the yeast strains, performed the experiments, and analyzed the data. NT and MDC contributed to the bioinformatics analysis of the sequencing data. DF and FS conceptualized the research project. EKP, CM, GL, AQ, and JMG contributed with technical support, resources, and intellectual support. DF, NT, and FS wrote the manuscript with insight from all the authors. The final version of the manuscript was approved by all the authors.

