## Supplementary information for "Optogenetic control of a horizontally acquired region in yeast prevent stuck fermentations"

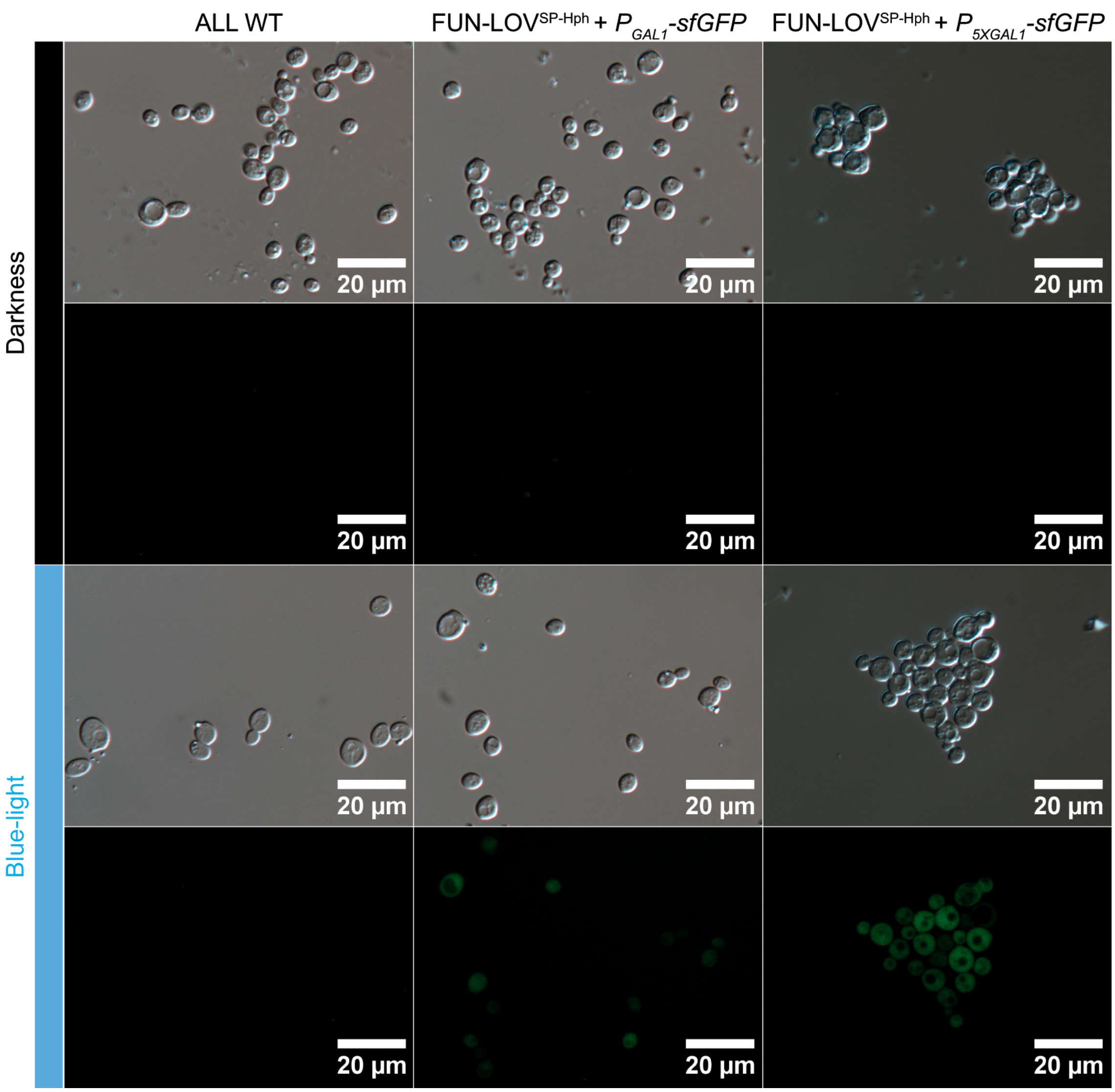


**Supplementary Figure S1.** Fluorescence microscopy for strains expressing *sfGFP* under the control of FUN-LOV$^{SP-Hph}$. *sfGFP* was expressed under the control of the FUN-LOV$^{SP-Hph}$ variant using either the $P_{GAL1}$ or $P_{5XGAL1}$ promoter. Fluorescence was evaluated in constant darkness (black bar, repression) or constant blue-light (blue bar, overexpression) conditions. Bright-field and fluorescence microscopy images are shown for each illumination condition. The scale bar represents 20 µm.

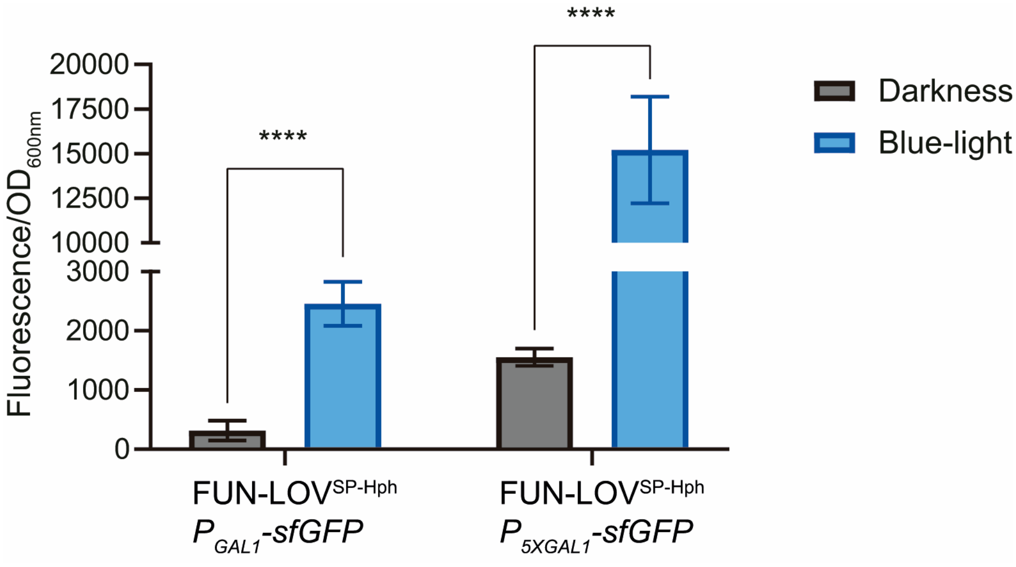


**Supplementary Figure S2.** Fluorescence measurements in yeast strains expressing *sfGFP* under the control of the FUN-LOV$^{SP\text{-}Hph}$ variant. *sfGFP* was expressed under the control of either the $P_{GAL1}$ or $P_{5XGAL1}$ promoter, which are recognized by FUN-LOV$^{SP\text{-}Hph}$. Fluorescence of the yeast cultures was evaluated in constant darkness (black bars) and constant blue-light (blue bars) conditions. The final fluorescence of the yeast cultures was normalized by $OD_{600nm}$. The average of six biological replicates with the standard deviation represented as error bars is shown. The asterisks represent a statistically significant difference between constant darkness and constant blue-light conditions (*t*-test, **** = $p < 0.0001$).

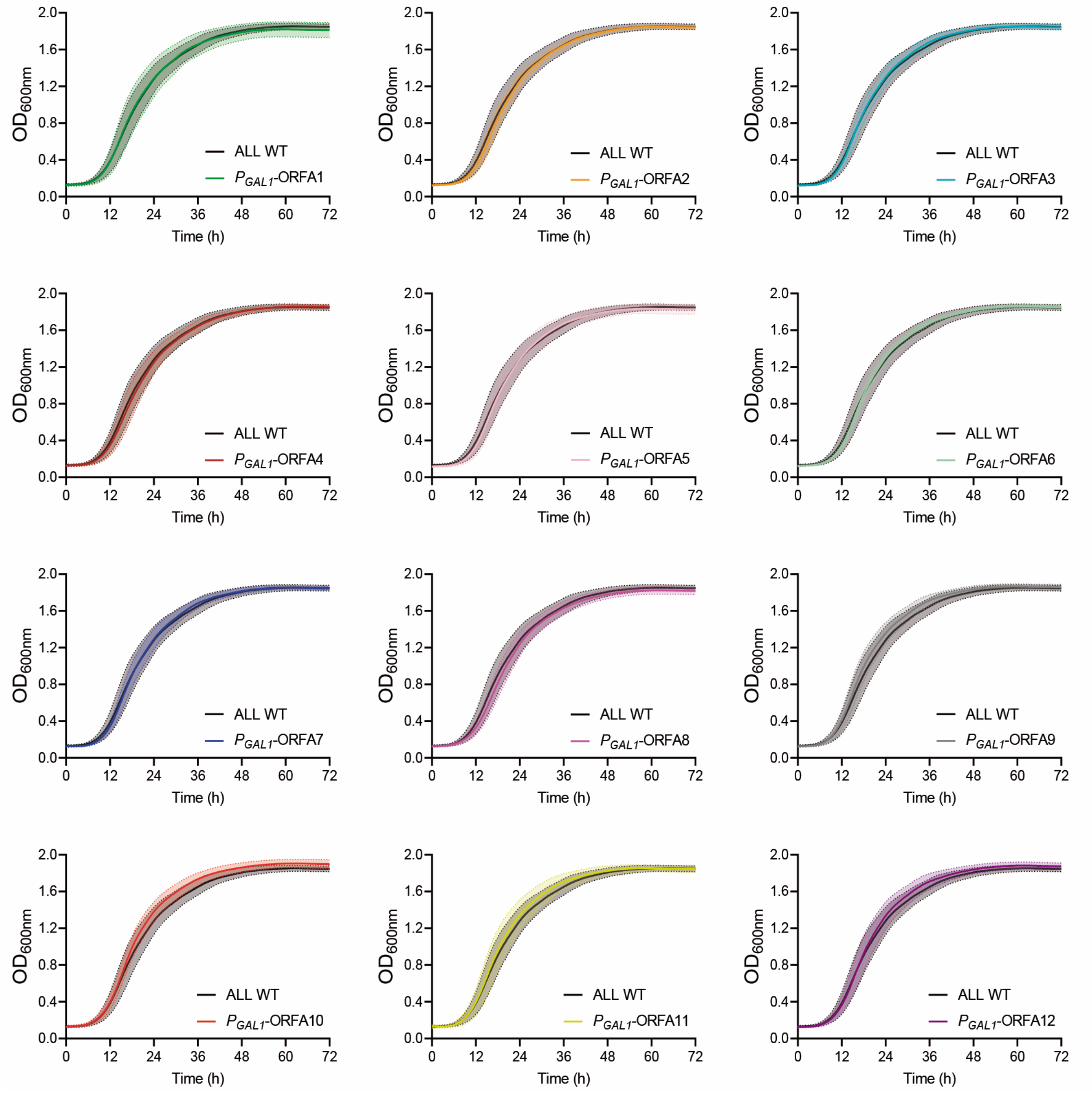


**Supplementary Figure S3.** Raw data for growth curves in SM300 and constant darkness conditions. The panels show the growth kinetics measured as Optical Density OD at 600 nm ($OD_{600nm}$) for the 'ALL' wild type strain and derived strains carrying the FUN-LOV$^{SP\text{-}Hph}$ variant controlling different ORFs within region A. The *GAL1* promoter ($P_{GAL1}$) is recognized by the FUN-LOV$^{SP\text{-}Hph}$ variant. In all panels, the average of six biological replicates with the standard deviation represented as a color shaded region is shown.

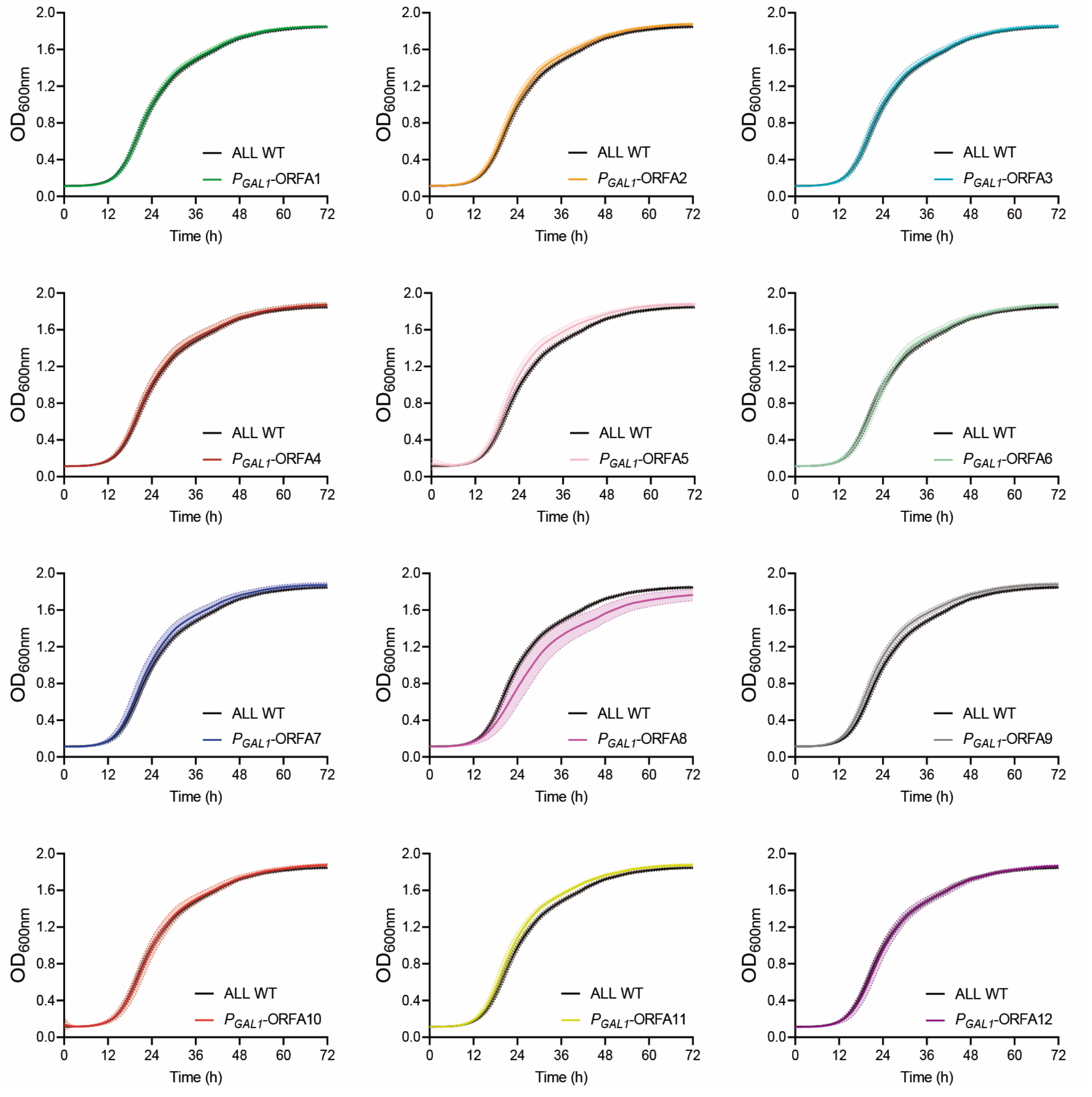


**Supplementary Figure S4.** Raw data for growth curves in SM300 and constant blue-light conditions. The panels show the growth kinetics measured as Optical Density OD at 600 nm ($OD_{600nm}$) for the 'ALL' wild type strain and derived strains carrying the FUN-LOV$^{SP\text{-}Hph}$ variant controlling different ORFs within region A. The *GAL1* promoter ($P_{GAL1}$) is recognized by the FUN-LOV$^{SP\text{-}Hph}$ variant. In all panels, the average of six biological replicates with the standard deviation represented as a color shaded region is shown.

.

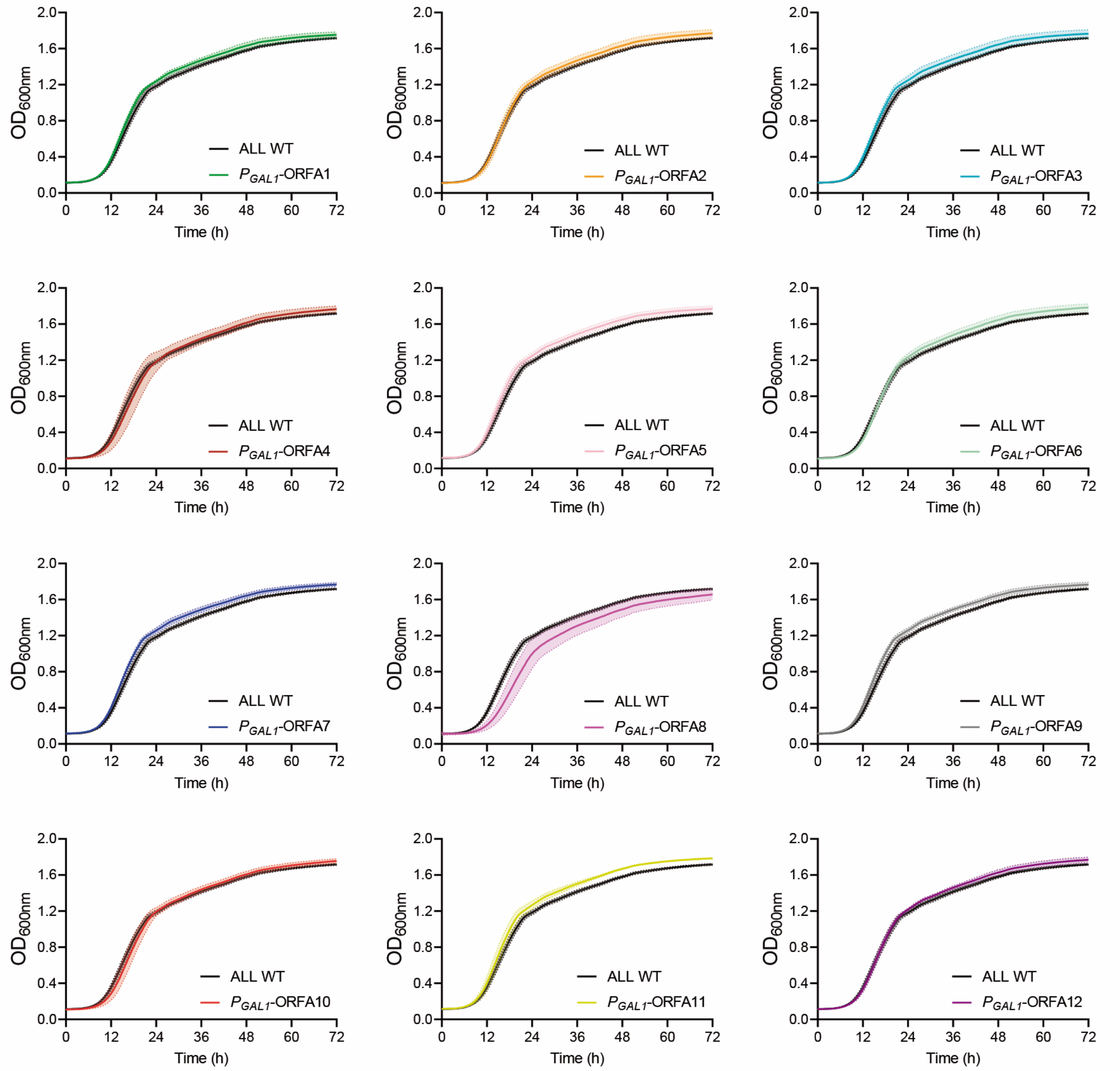


**Supplementary Figure S5.** Raw data for growth curves in SM140 and constant darkness conditions. The panels show the growth kinetics measured as Optical Density OD at 600 nm ($OD_{600nm}$) for the 'ALL' wild type strain and derived strains carrying the FUN-LOV$^{SP\text{-}Hph}$ variant controlling different ORFs within region A. The *GAL1* promoter ($P_{GAL1}$) is recognized by the FUN-LOV$^{SP\text{-}Hph}$ variant. In all panels, the average of six biological replicates with the standard deviation represented as a color shaded region is shown.

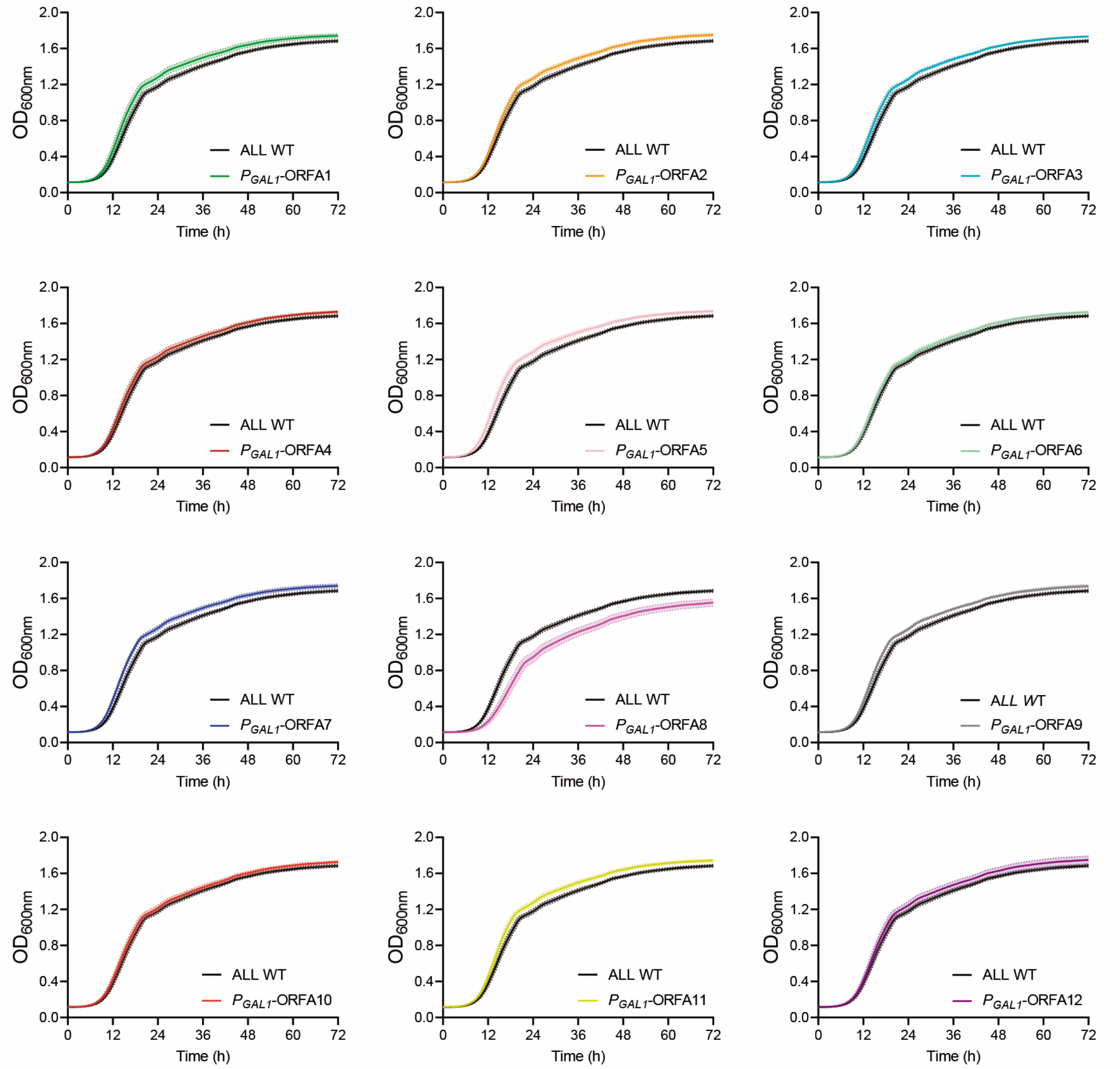


**Supplementary Figure S6.** Raw data for growth curves in SM140 and constant blue-light conditions. The panels show the growth kinetics measured as Optical Density OD at 600 nm ($OD_{600nm}$) for the 'ALL' wild type strain and derived strains carrying the FUN-LOV$^{SP\text{-}Hph}$ variant controlling different ORFs within region A. The *GAL1* promoter ($P_{GAL1}$) is recognized by the FUN-LOV$^{SP\text{-}Hph}$ variant. In all panels, the average of six biological replicates with the standard deviation represented as a color shaded region is shown.

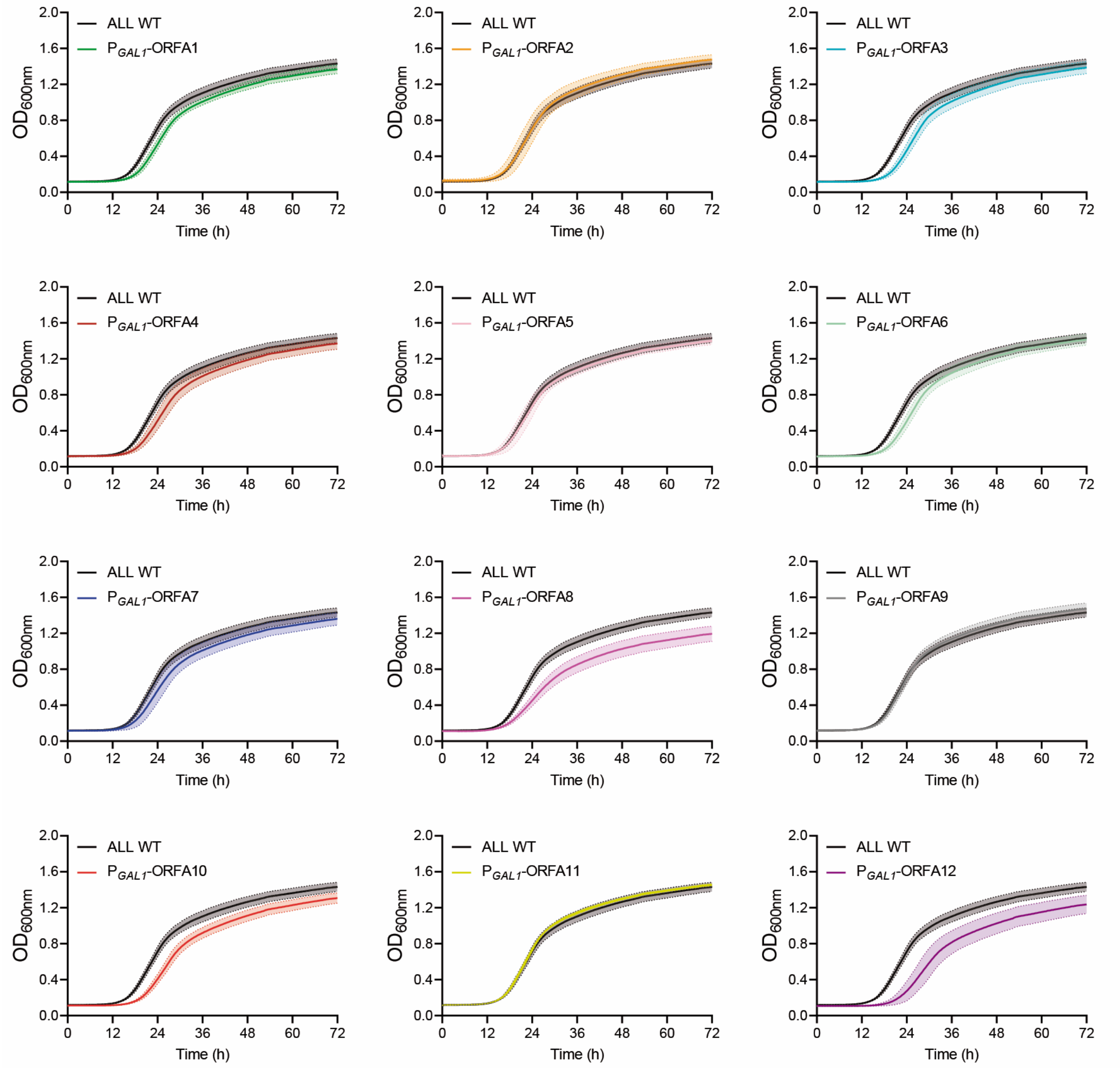


**Supplementary Figure S7.** Raw data for growth curves in SM60 and constant darkness conditions. The panels show the growth kinetics measured as Optical Density OD at 600 nm ($OD_{600nm}$) for the 'ALL' wild type strain and derived strains carrying the FUN-LOV$^{SP\text{-}Hph}$ variant controlling different ORFs within region A. The *GAL1* promoter ($P_{GAL1}$) is recognized by the FUN-LOV$^{SP\text{-}Hph}$ variant. In all panels, the average of six biological replicates with the standard deviation represented as a color shaded region is shown.

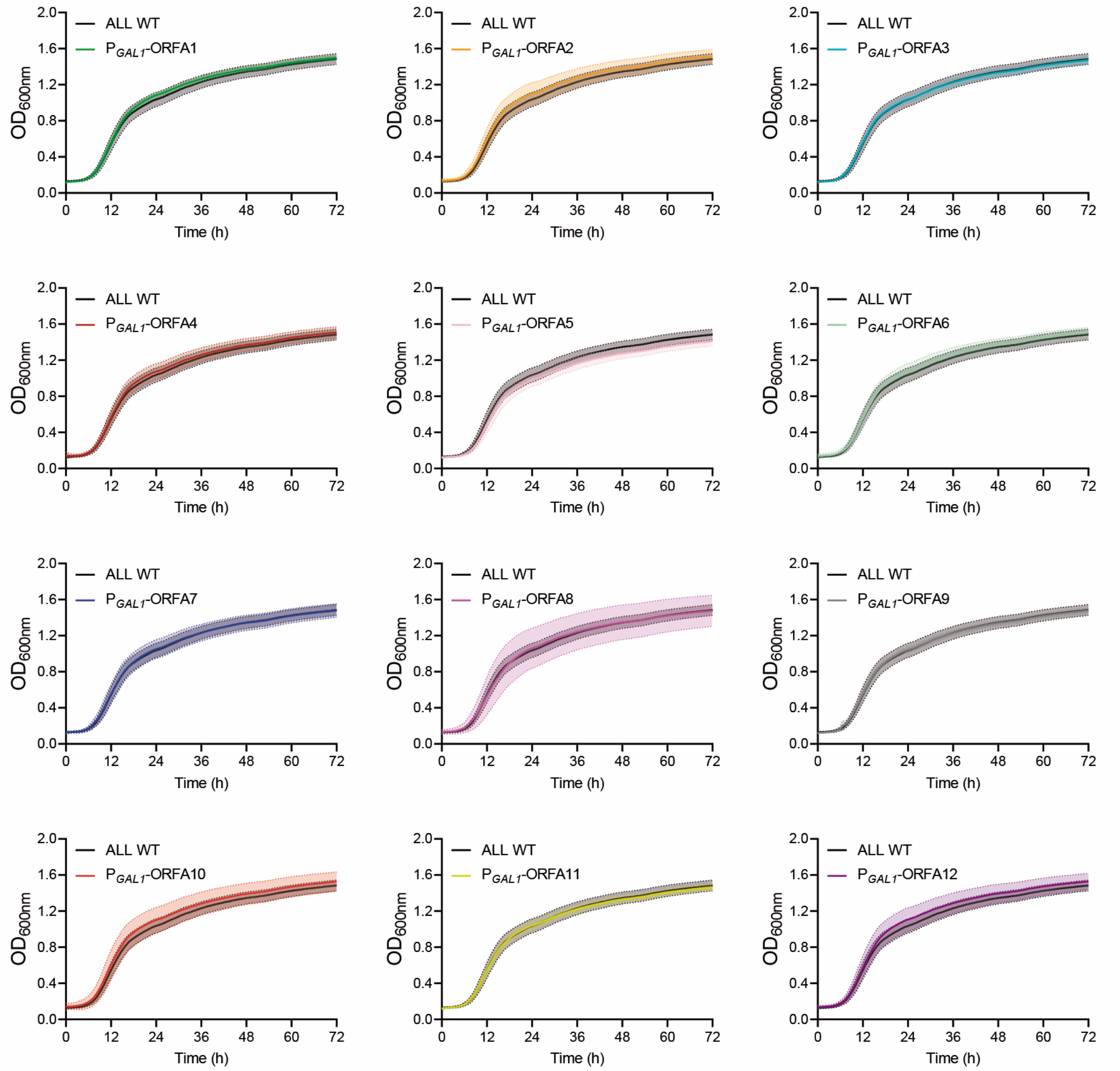


**Supplementary Figure S8.** Raw data for growth curves in SM60 and constant blue-light conditions. The panels show the growth kinetics measured as Optical Density OD at 600 nm ($OD_{600nm}$) for the 'ALL' wild type strain and derived strains carrying the FUN-LOV$^{SP\text{-}Hph}$ variant controlling different ORFs within region A. The *GAL1* promoter ($P_{GAL1}$) is recognized by the FUN-LOV$^{SP\text{-}Hph}$ variant. In all panels, the average of six biological replicates with the standard deviation represented as a color shaded region is shown.

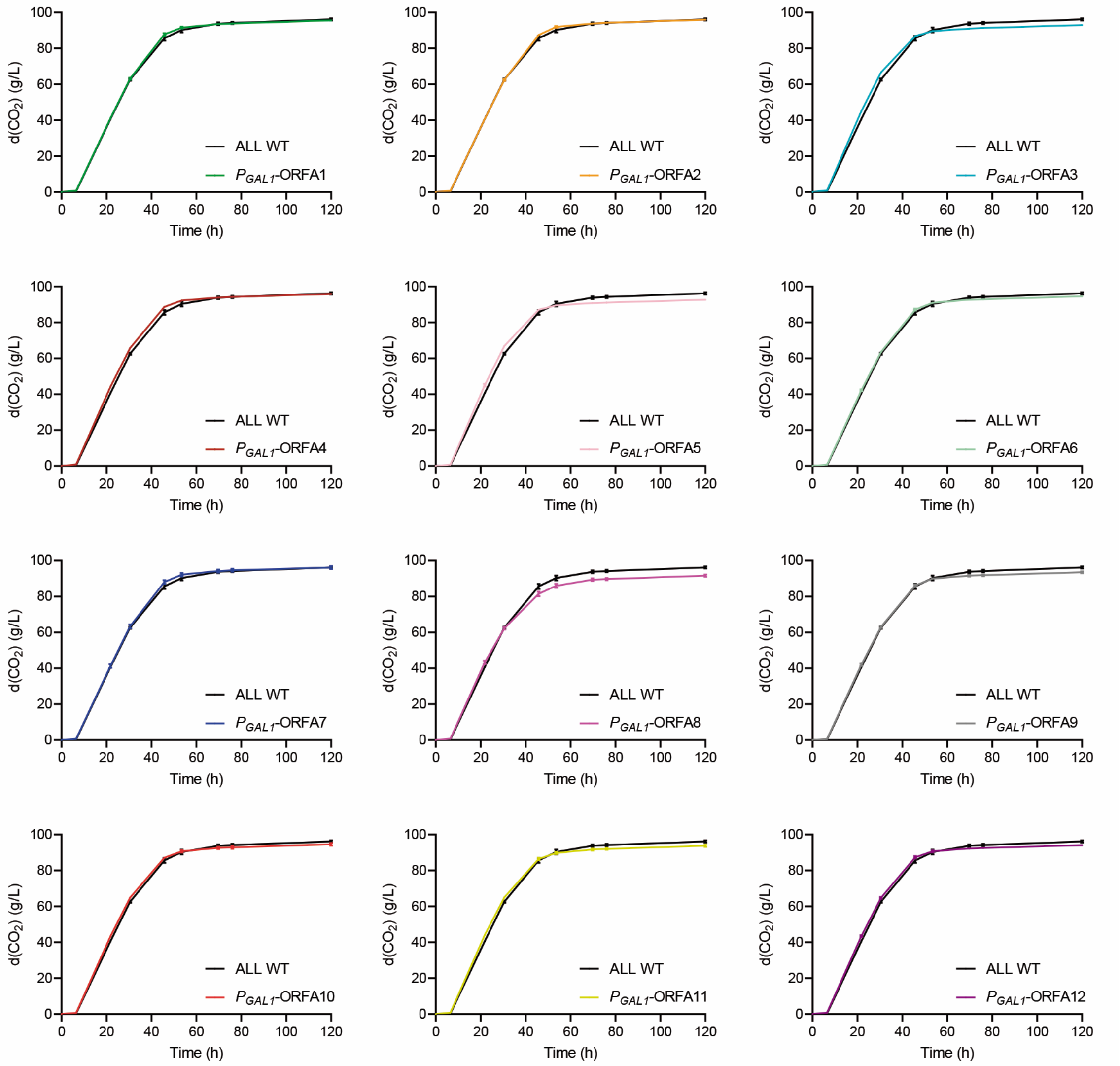


**Supplementary Figure S9.** Raw data for fermentations in SM300 and constant darkness conditions. The panels show the fermentation kinetics measured as $CO_2$ loss for the 'ALL' wild type strain and derived versions carrying the FUN-LOV$^{SP\text{-}Hph}$ variant controlling different ORFs within region A. The *GAL1* promoter ($P_{GAL1}$) is recognized by the FUN-LOV$^{SP\text{-}Hph}$ variant. In all panels, the average of three biological replicates with the standard deviation represented as error bars is shown.

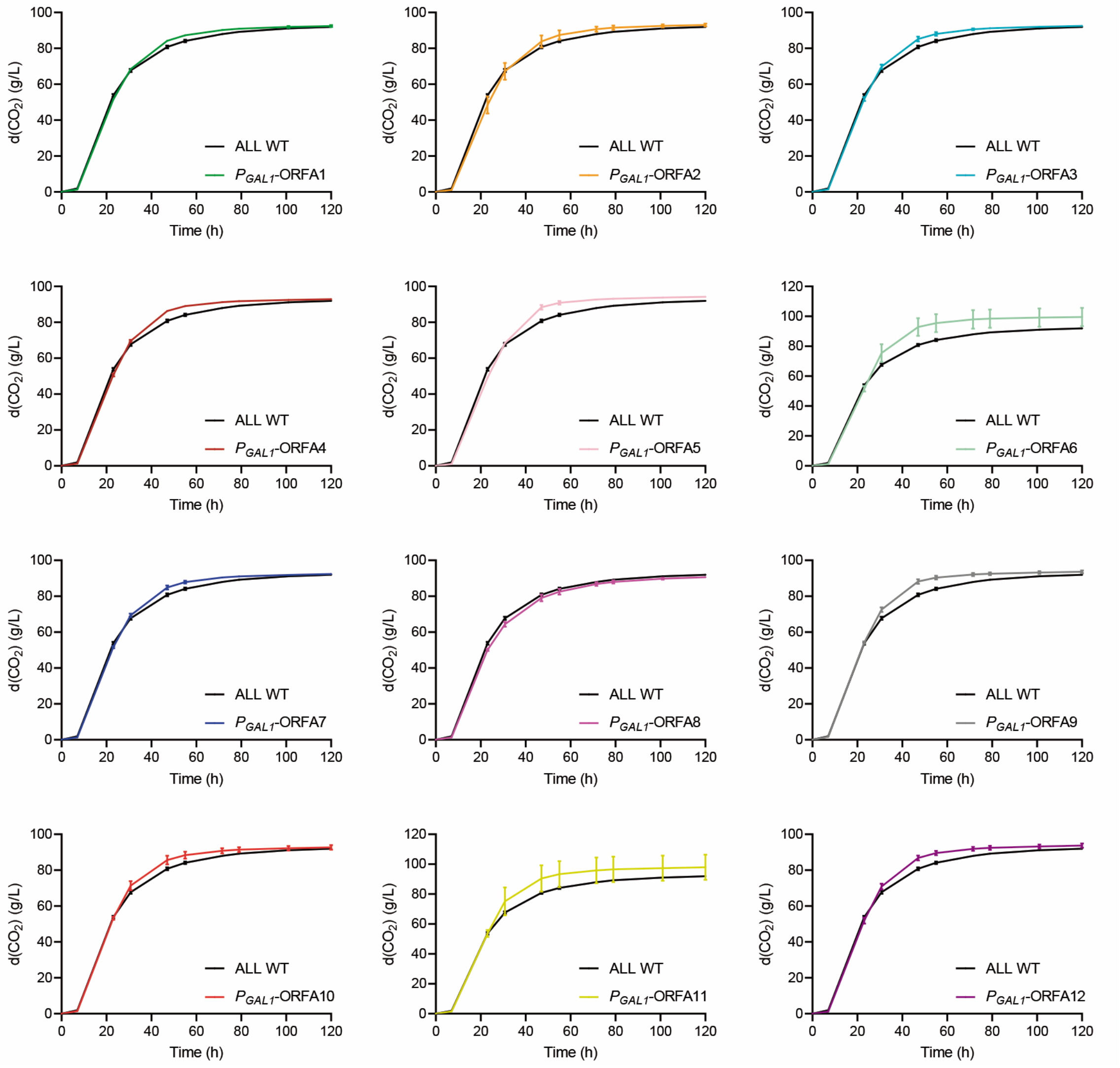


**Supplementary Figure S10.** Raw data for fermentations in SM300 and constant blue-light conditions. The panels show the fermentation kinetics measured as $CO_2$ loss for the 'ALL' wild type strain and derived versions carrying the FUN-LOV$^{SP\text{-}Hph}$ variant controlling different ORFs within region A. The *GAL1* promoter ($P_{GAL1}$) is recognized by the FUN-LOV$^{SP\text{-}Hph}$ variant. In all panels, the average of three biological replicates with the standard deviation represented as error bars is shown.

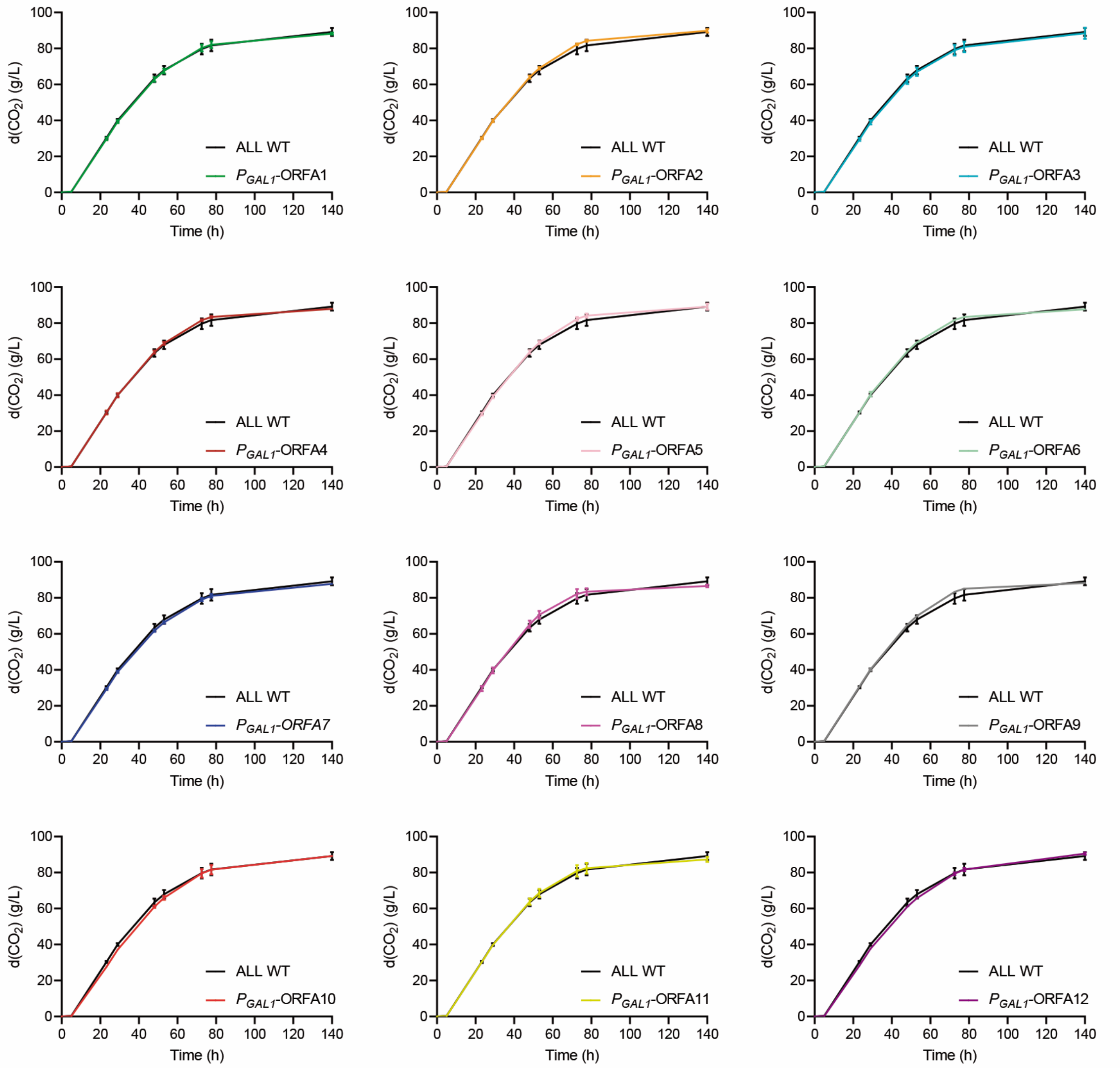


**Supplementary Figure S11.** Raw data for fermentations in SM140 and constant darkness conditions. The panels show the fermentation kinetics measured as $CO_2$ loss for the 'ALL' wild type strain and derived versions carrying the FUN-LOV$^{SP\text{-}Hph}$ variant controlling different ORFs within region A. The *GAL1* promoter ($P_{GAL1}$) is recognized by the FUN-LOV$^{SP\text{-}Hph}$ variant. In all panels, the average of three biological replicates with the standard deviation represented as error bars is shown.

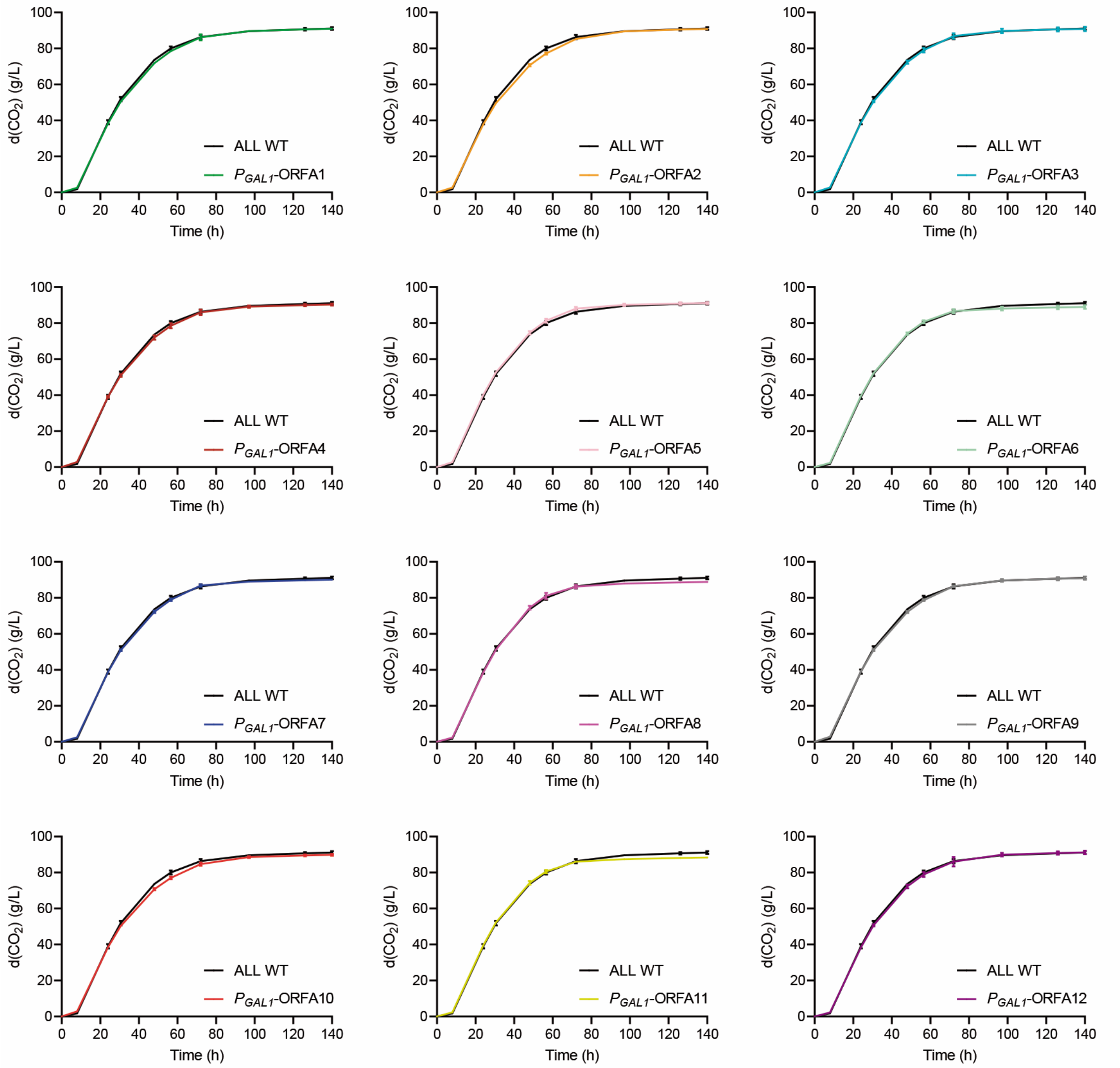


**Supplementary Figure S12.** Raw data for fermentations in SM140 and constant blue-light conditions. The panels show the fermentation kinetics measured as $CO_2$ loss for the 'ALL' wild type strain and derived versions carrying the FUN-LOV$^{SP\text{-}Hph}$ variant controlling different ORFs within region A. The *GAL1* promoter ($P_{GAL1}$) is recognized by the FUN-LOV$^{SP\text{-}Hph}$ variant. In all panels, the average of three biological replicates with the standard deviation represented as error bars is shown.

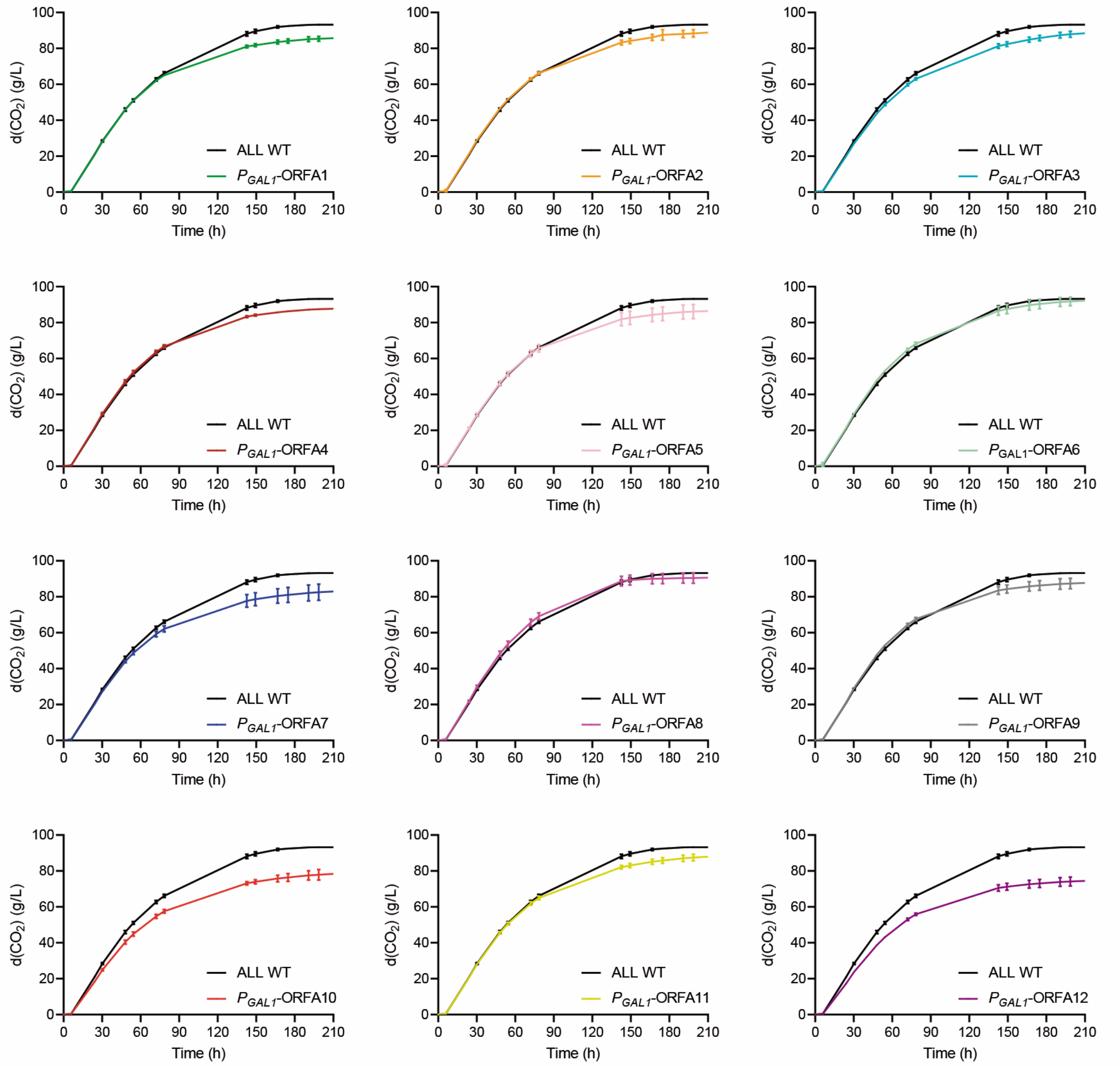


**Supplementary Figure S13.** Raw data for fermentations in SM60 and constant darkness conditions. The panels show the fermentation kinetics measured as $CO_2$ loss for the 'ALL' wild type strain and derived versions carrying the FUN-LOV$^{SP\text{-}Hph}$ variant controlling different ORFs within region A. The *GAL1* promoter ($P_{GAL1}$) is recognized by the FUN-LOV$^{SP\text{-}Hph}$ variant. In all panels, the average of three biological replicates with the standard deviation represented as error bars is shown.

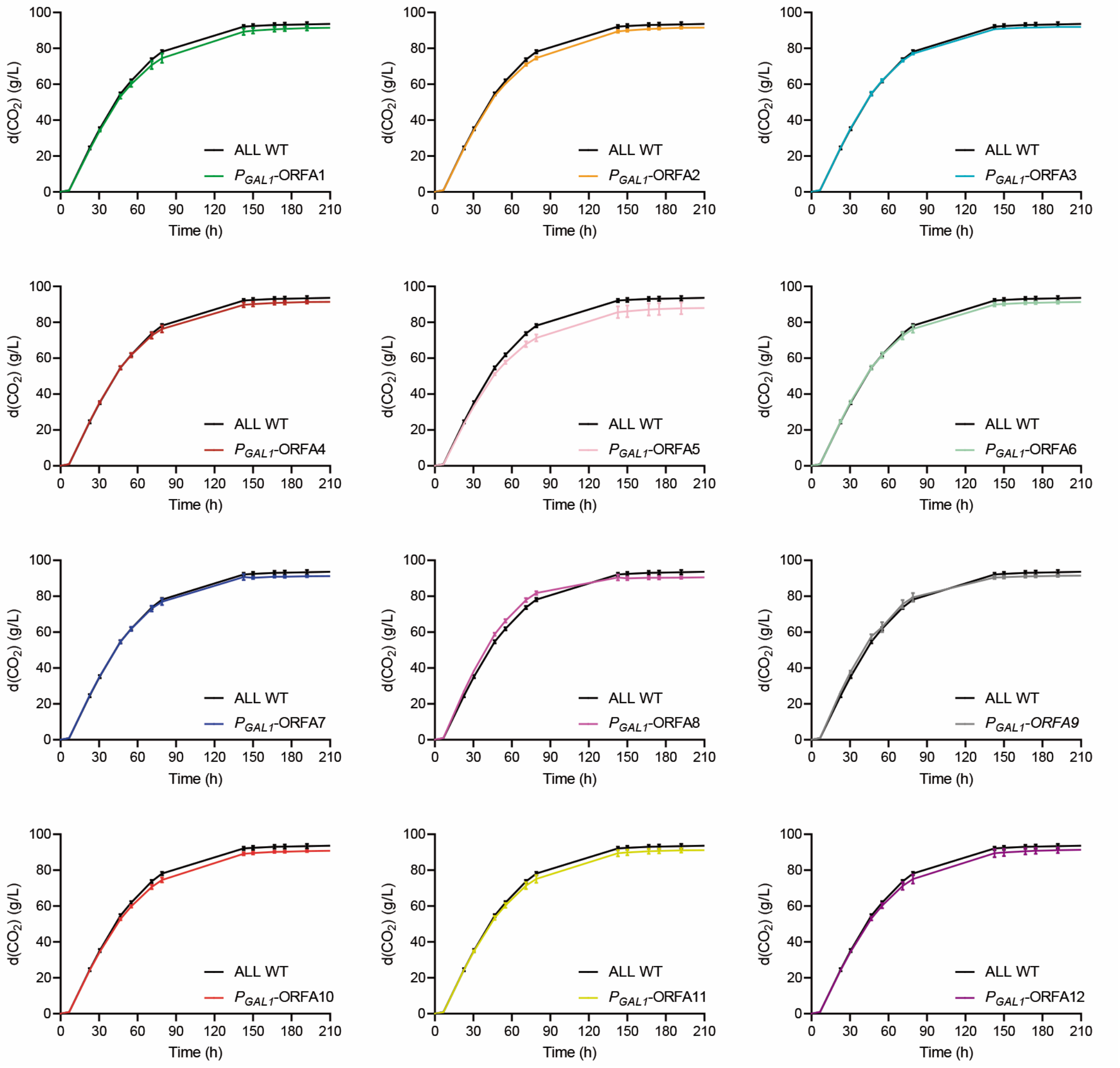


**Supplementary Figure S14.** Raw data for fermentations in SM60 and constant blue-light conditions. The panels show the fermentation kinetics measured as $CO_2$ loss for the 'ALL' wild type strain and the derived versions carrying the FUN-LOV$^{SP\text{-}Hph}$ variant controlling different ORFs within region A. The *GAL1* promoter ($P_{GAL1}$) is recognized by the FUN-LOV$^{SP\text{-}Hph}$ variant. In all panels, the average of three biological replicates with the standard deviation represented as error bars is shown.

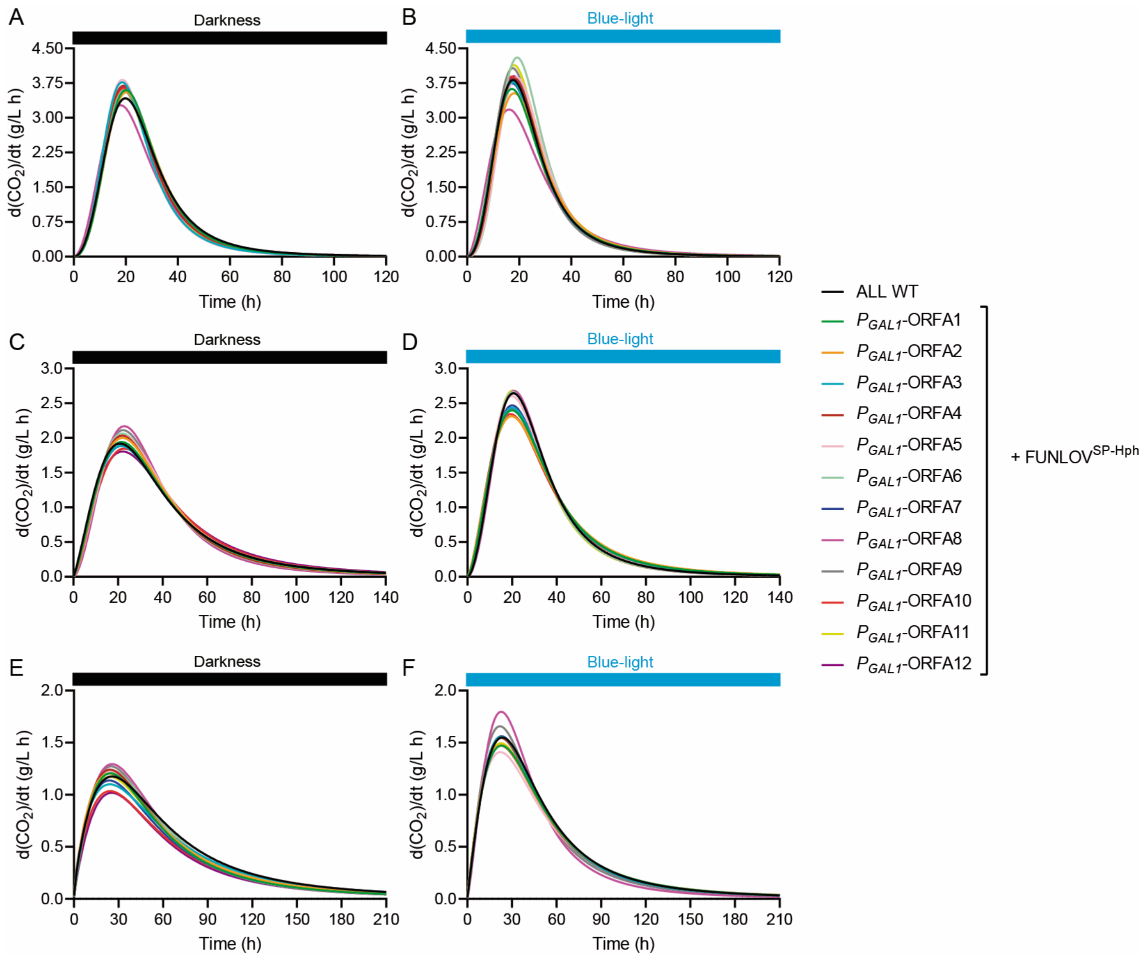


**Supplementary Figure S15.** Maximal $CO_2$ production rate ($V_{max}$) derived from fermentation kinetics. The 'ALL' wine yeast strain and derived strains carrying the FUN-LOV$^{SP\text{-}Hph}$ variant controlling the expression of different ORFs within region A were subjected to fermentations in SM300 (panels A and B), SM140 (panels C and D), and SM60 (panels E and F). Fermentations were performed in constant darkness (panels A, C, and E) and constant blue-light (panels B, D, and F) conditions. The $CO_2$ release curves were fitted to a sigmoid non-linear regression, and the first derivate was calculated to obtain the maximal $CO_2$ production rate ($V_{max}$, peak of the curves). The average of three biological replicates is shown.

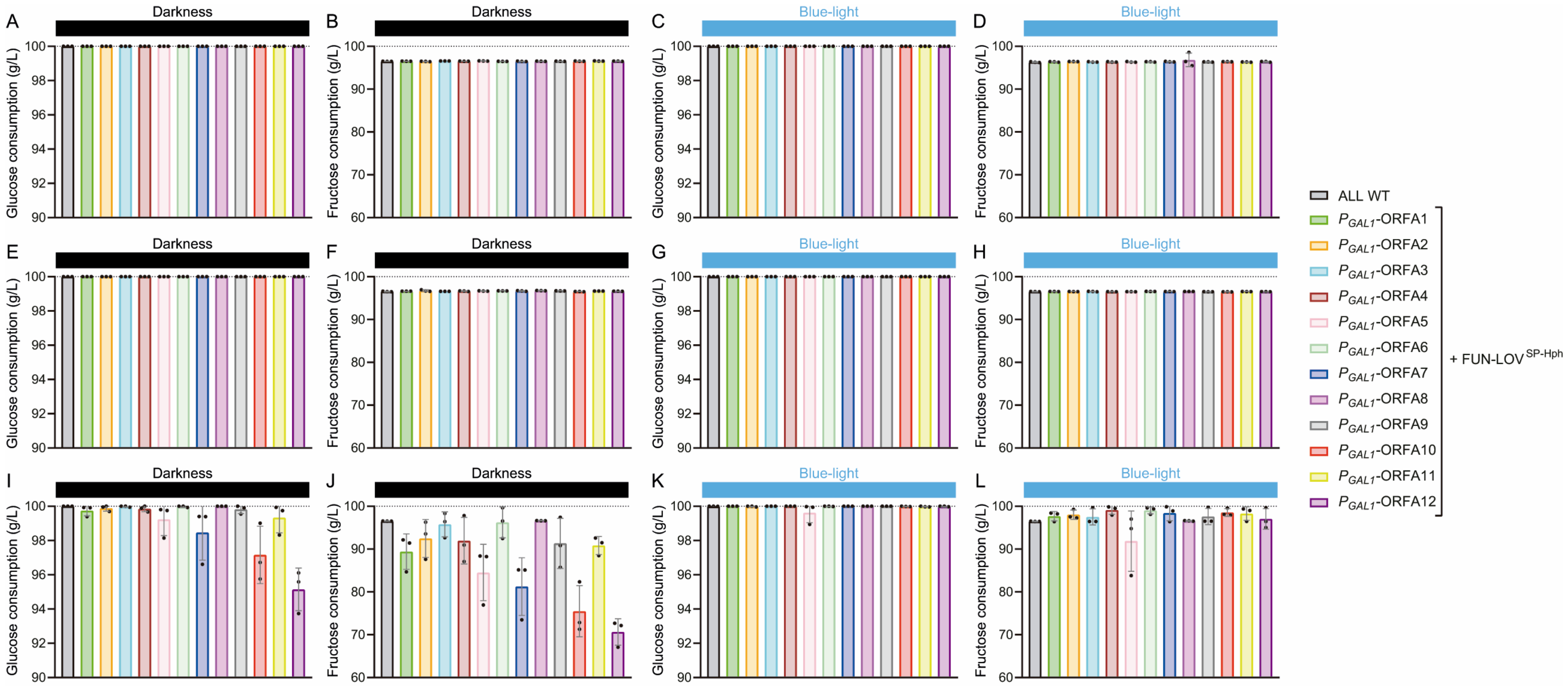


**Supplementary Figure S16.** Sugar consumption at the end of the fermentations. Fermentations were performed in SM300 (panels A, B, C, and D), SM140 (panels E, F, G, and H), and SM60 (panels I, J, K, and L) for the 'ALL' wine yeast strain and derived strains carrying the FUN-LOV$^{SP-Hph}$ variant controlling the expression of different ORFs within region A. Fermentations were performed in constant darkness (panels A, B, E, F, I, and J) and constant blue-light (panels C, D, G, H, K, and L) conditions. The fermentation medium was supplemented with 100 g/L of glucose and 100 g/L of fructose (dashed lines) and consumption were determined by HPLC. In all panels, the average of three biological replicates with the standard deviation represented as error bars is shown.

**Supplementary Table S1.** Yeast strains used and generated in this work.

| Strain | Genotype | Source |
|---|---|---|
| ALL WT | - | Peter et al., 2018 |
| $P_{GAL1}$-*Luc* + FUN-LOV$^{SP\text{-}Hph}$ | ALL; *gal3*Δ::KanMxRV $P_{GAL1}$-*Luc*; *ho*Δ::FUN-LOV$^{SP\text{-}Hph}$ | This work |
| $P_{5XGAL1}$-*Luc* + FUN-LOV$^{SP\text{-}Hph}$ | ALL; *gal3*Δ::KanMxRV $P_{5XGAL1}$-*Luc*; *ho*Δ::FUN-LOV$^{SP\text{-}Hph}$ | This work |
| $P_{GAL1}$-*sfGFP* + FUN-LOV$^{SP\text{-}Hph}$ | ALL; *gal3*Δ::KanMxRV $P_{GAL1}$-*sfGFP*; *ho*Δ::FUN-LOV$^{SP\text{-}Hph}$ | This work |
| $P_{5XGAL1}$-*sfGFP* + FUN-LOV$^{SP\text{-}Hph}$ | ALL; *gal3*Δ::KanMxRV $P_{5XGAL1}$-*sfGFP*; *ho*Δ::FUN-LOV$^{SP\text{-}Hph}$ | This work |
| $P_{GAL1}$-ORFA1 + FUN-LOV$^{SP\text{-}Hph}$ | ALL; $P_{ORFA1}$Δ::KanMxRV-$P_{GAL1}$; *ho*Δ::FUN-LOV$^{SP\text{-}Hph}$ | This work |
| $P_{GAL1}$-ORFA2 + FUN-LOV$^{SP\text{-}Hph}$ | ALL; $P_{ORFA2}$Δ::KanMxRV-$P_{GAL1}$; *ho*Δ::FUN-LOV$^{SP\text{-}Hph}$ | This work |
| $P_{GAL1}$-ORFA3 + FUN-LOV$^{SP\text{-}Hph}$ | ALL; $P_{ORFA3}$Δ::KanMxRV-$P_{GAL1}$; *ho*Δ::FUN-LOV$^{SP\text{-}Hph}$ | This work |
| $P_{GAL1}$-ORFA4 + FUN-LOV$^{SP\text{-}Hph}$ | ALL; $P_{ORFA4}$Δ::KanMxRV-$P_{GAL1}$; *ho*Δ::FUN-LOV$^{SP\text{-}Hph}$ | This work |
| $P_{GAL1}$-ORFA5 + FUN-LOV$^{SP\text{-}Hph}$ | ALL; $P_{ORFA5}$Δ::KanMxRV-$P_{GAL1}$; *ho*Δ::FUN-LOV$^{SP\text{-}Hph}$ | This work |
| $P_{GAL1}$-ORFA6 + FUN-LOV$^{SP\text{-}Hph}$ | ALL; $P_{ORFA6}$Δ::KanMxRV-$P_{GAL1}$; *ho*Δ::FUN-LOV$^{SP\text{-}Hph}$ | This work |
| $P_{GAL1}$-ORFA7 + FUN-LOV$^{SP\text{-}Hph}$ | ALL; $P_{ORFA7}$Δ::KanMxRV-$P_{GAL1}$; *ho*Δ::FUN-LOV$^{SP\text{-}Hph}$ | This work |
| $P_{GAL1}$-ORFA8 + FUN-LOV$^{SP\text{-}Hph}$ | ALL; $P_{ORFA8}$Δ::KanMxRV-$P_{GAL1}$; *ho*Δ::FUN-LOV$^{SP\text{-}Hph}$ | This work |
| $P_{GAL1}$-ORFA9 + FUN-LOV$^{SP\text{-}Hph}$ | ALL; $P_{ORFA9}$Δ::KanMxRV-$P_{GAL1}$; *ho*Δ::FUN-LOV$^{SP\text{-}Hph}$ | This work |
| $P_{GAL1}$-ORFA10 + FUN-LOV$^{SP\text{-}Hph}$ | ALL; $P_{ORFA10}$Δ::KanMxRV-$P_{GAL1}$; *ho*Δ::FUN-LOV$^{SP\text{-}Hph}$ | This work |
| $P_{GAL1}$-ORFA11 + FUN-LOV$^{SP\text{-}Hph}$ | ALL; $P_{ORFA11}$Δ::KanMxRV-$P_{GAL1}$; *ho*Δ::FUN-LOV$^{SP\text{-}Hph}$ | This work |
| $P_{GAL1}$-ORFA12 + FUN-LOV$^{SP\text{-}Hph}$ | ALL; $P_{ORFA12}$Δ::KanMxRV-$P_{GAL1}$; *ho*Δ::FUN-LOV$^{SP\text{-}Hph}$ | This work |
| $P_{5XGAL1}$-ORFA6 + FUN-LOV$^{SP\text{-}Hph}$ | ALL; $P_{ORFA6}$Δ::KanMxRV-$P_{5XGAL1}$; *ho*Δ::FUN-LOV$^{SP\text{-}Hph}$ | This work |
| $P_{5XGAL1}$-ORFA8 + FUN-LOV$^{SP\text{-}Hph}$ | ALL; $P_{ORFA8}$Δ::KanMxRV-$P_{5XGAL1}$; *ho*Δ::FUN-LOV$^{SP\text{-}Hph}$ | This work |

**Supplementary Table S2.** List of primers used and generated in this work.

| Description | Type | Length (nt) | Sequence (5´-3´) |
|---|---|---|---|
| Swapping *GAL3* locus | Fw | 70 | AGGAGTGCAAAAAGAGAAAATAAAAGTAAAAAGGTAGG GCAACACATAGTATCGATGAATTCGAGCTCGT |
| Swapping *GAL3* locus | Rv | 70 | TATGAGTAAACTTTTAATATTTAAAGGTTGTTCCAAGAAGG TGTTTAGTGTGGATCCTTGCAAATTAAAG |
| *GAL3* Upstream | Fw | 20 | ATGAAATCGCCATGCCAAGC |
| *GAL3* Downstream | Rv | 20 | GTGCGGAGCCACTCTGACTC |
| K3: internal KanMx | Fw | 21 | CATCCTATGGAACTGCCTCGG |
| Internal *Luc* | Fw | 20 | ATCGTGGTGTGCTCTGAGAA |
| Assemble pRS316-KanMx | Fw | 60 | GGCCAGTGAATTGTAATACGACTCACTATAGGGCGAATTG ATCGATGAATTCGAGCTCGT |
| Assemble $P_{GAL1/5XGAL1}$-*sfGFP* | Fw | 60 | AGCTGTAATACGACTCACTATAGGGAATATTAAGCTTACCA TGCGTAAAGGCGAAGAGCT |
| Assemble $P_{GAL1/5XGAL1}$-*sfGFP* | Rv | 60 | TAGGGACGACACCAGTGAACAGCTCTTCGCCTTTACGCAT GGTAAGCTTAATATTCCCTA |
| Assemble *sfGFP-CYC1*$_{ter}$ | Fw | 60 | AGCGGGCATCACGCATGGTATGGATGAACTGTACAAATGA TCATGTAATTAGTTATGTCA |
| Assemble *sfGFP-CYC1*$_{ter}$ | Rv | 60 | GGGGAGGGCGTGAATGTAAGTGACATAACTAATTACATGA TCATTTGTACAGTTCATCCA |
| Assemble *CYC1*$_{ter}$-pRS316 | Rv | 60 | CAAGCTCGGAATTAACCCTCACTAAAGGGAACAAAAGCT GTGGATCCTTGCAAATTAAAG |
| pRS316 Upstream | Fw | 20 | TTCGCTATTACGCCAGCTGG |
| pRS316 Downstream | Rv | 20 | TGCTTCCGGCTCCTATGTTG |
| Swapping *HO* locus | Fw | 70 | TCTAAATCCATATCCTCATAAGCAGCAATCAATTCTATCTAT ACTTTAAAATCGATGAATTCGAGCTCGT |
| Swapping *HO* locus | Rv | 70 | ATTAAATTTTACTTTTATTACATACAACTTTTTAAACTAATA TACACATTTGGATCCTTGCAAATTAAAG |
| *HO* Upstream | Fw | 20 | GAATTGTACTACCGCTGGGC |
| *HO* Downstream | Rv | 22 | TGGTTGAAACAAATCAGTGCCG |
| Swapping $P_{ORFA1}$ | Fw | 70 | CCCAAACCATTATTTGCAGTTGATGCACGCTTCCCGTTCTC AGTTGTCATGGTAAGCTTAATATTCCCTA |
| Swapping $P_{ORFA1}$ | Rv | 70 | GGTTCGCTATTTGAGTACGTGGACTTGGGTCTGCTTGAAA CGCTACGATGATCGATGAATTCGAGCTCGT |
| Swapping $P_{ORFA1}$ confirmation | Rv | 20 | CCAGGTGCTGCTTCACGGGT |
| Swapping $P_{ORFA2}$ | Fw | 70 | ACTCGGAGTATTTTATTTATTGTCTAGCCTTTAATTGCGGAT GCGCCTTCATCGATGAATTCGAGCTCGT |
| Swapping $P_{ORFA2}$ | Rv | 70 | GACGATTCTCCTATGTAATGATGAAAATCAGGGTATGATCT TTTTAACATGGTAAGCTTAATATTCCCTA |
| Swapping $P_{ORFA2}$ confirmation | Fw | 20 | CGCGGACTTGTCCTTGGGGT |

| | | | |
|---|---|---|---|
| Swapping $P_{ORFA3}$ | Fw | 70 | ATACTAGATTTAGATTGTCTTTACATTAATAATCTAGATTCTTCAGTTTCATCGATGAATTCGAGCTCGT |
| Swapping $P_{ORFA3}$ | Rv | 70 | ACTTCATCAATTGCGATGGCTTTTTCGTCGAGGTTTTTGGTTTCAGACATGGTAAGCTTAATATTCCCTA |
| Swapping $P_{ORFA3}$ confirmation | Fw | 20 | AGACTGGTATGCGTCGTTGG |
| Swapping $P_{ORFA4}$ | Fw | 70 | ATATGTTGAGCTATAAATCCATTTGCACCCGAAACCAACACAGCAGACATGGTAAGCTTAATATTCCCTA |
| Swapping $P_{ORFA4}$ | Rv | 70 | ACGCGCACCCTTACATTTCTCGAAGTTCTTCAAAGCTTCAGACAGGGCCAATCGATGAATTCGAGCTCGT |
| Swapping $P_{ORFA4}$ confirmation | Rv | 20 | GGGGGCCAAACAGCGGATGT |
| Swapping $P_{ORFA5}$ | Fw | 70 | CCAGTCACCAGCACCCATTTGCCTGATATTTTTCTTTCTGTCGTTCTCATGGTAAGCTTAATATTCCCTA |
| Swapping $P_{ORFA5}$ | Rv | 70 | GGCACCAGTGTAATTGTGGTTGAGGAGGATTACACCTCCAAGACCTGCTCATCGATGAATTCGAGCTCGT |
| Swapping $P_{ORFA5}$ confirmation | Rv | 20 | CGTGCGCACGGAAAAGCTGA |
| Swapping $P_{ORFA6}$ | Fw | 70 | CAGCCGTTGTGGCTATATCTTTTACGCGGCGTACCAGTTACTCTGGTCATGGTAAGCTTAATATTCCCTA |
| Swapping $P_{ORFA6}$ | Rv | 70 | ACTTTCAAGTATATACGTCCTCAGTTTATTTCCAGACATACTTATATAAAATCGATGAATTCGAGCTCGT |
| Swapping $P_{ORFA6}$ confirmation | Rv | 20 | GAGATCACCTCAGTCGTCGC |
| Swapping $P_{ORFA7}$ | Fw | 70 | CCATCCAAGAAATTGCCAAAATAGGCATCCAAAAATGACTTGTTGACCATGGTAAGCTTAATATTCCCTA |
| Swapping $P_{ORFA7}$ | Rv | 70 | TGGTATGAATGAACTGGGCTATATTGTATCCCAAGAGCCGAAGTTGACATATCGATGAATTCGAGCTCGT |
| Swapping $P_{ORFA7}$ confirmation | Rv | 20 | TGCCGATGTGCCCGACAGAG |
| Swapping $P_{ORFA8}$ | Fw | 70 | TGCTTCGGTGTGAGCAGAAGCTTGATGATTCCGATCTCCATCGCTGCGTTATCGATGAATTCGAGCTCGT |
| Swapping $P_{ORFA8}$ | Rv | 70 | AGTTTCGATTTGTGATCAGGTAATACCTGTAAAACGGGGTCATTTTTCATGGTAAGCTTAATATTCCCTA |
| Swapping $P_{ORFA8}$ confirmation | Fw | 20 | CCCTTGACAGATACAGCCCC |
| Swapping $P_{ORFA9}$ | Fw | 70 | ACTTGCAAAAATTTGTCGCATCTTTTGAGCGTTTCGATGTTAAAAAGCATGGTAAGCTTAATATTCCCTA |
| Swapping $P_{ORFA9}$ | Rv | 70 | GGAGGTAGAAGAACGGAAGACGCACCACTTGTCATCAAATGCTCACAACAATCGATGAATTCGAGCTCGT |
| Swapping $P_{ORFA9}$ confirmation | Rv | 20 | TCGCCCGGGGGTAGTTCTCA |
| Swapping $P_{ORFA10}$ | Fw | 70 | TAATCCTTATAATACTGAGCCCTCGCATCCCAAAAAGCGTTCCCGTCCATGGTAAGCTTAATATTCCCTA |
| Swapping $P_{ORFA10}$ | Rv | 70 | TTTTGAGAATGCTACAGATCAAAGCTTTAATGTTGATAGAGATAAATAGTATCGATGAATTCGAGCTCGT |
| Swapping $P_{ORFA10}$ confirmation | Rv | 20 | CGGTCATGTCGGTGATGGCA |

| | | | |
|---|---|---|---|
| Swapping $P_{ORFA11}$ | Fw | 70 | TTTGGATAGGCCTCAGCAGTTAACTGAGCAACAGGAGTTT GAATGGTCATGGTAAGCTTAATATTCCCTA |
| Swapping $P_{ORFA11}$ | Rv | 70 | GATCCCCTTTCTTCAAGGACTATGGGAAAGAAAGAATGCT ACAGTATTTGATCGATGAATTCGAGCTCGT |
| Swapping $P_{ORFA11}$ confirmation | Rv | 20 | TGCACTTTGGCACCGCCTCA |
| Swapping $P_{ORFA12}$ | Fw | 70 | CCTGCCATAGGAGCCTGGATGATTGGGTATTTCAAACTCA GACGAGTCATGGTAAGCTTAATATTCCCTA |
| Swapping $P_{ORFA12}$ | Rv | 70 | ATCATCTTGAGTCGCACGATGCTGAAGTCAGTAATCGAAG TTTGTCTTGAATCGATGAATTCGAGCTCGT |
| Swapping $P_{ORFA12}$ confirmation | Rv | 20 | GATCGTTGACCACGCGGCGT |
| ORFA6 qPCR | Fw | 20 | GCATGATTCTTTCACGGCGT |
| ORFA6 qPCR | Rv | 20 | GGCTCCTCAATAGCCGTCAT |
| ORFA8 qPCR | Fw | 20 | ATGCCACTGTTTCTGTCGGA |
| ORFA8 qPCR | Rv | 20 | GCCAATATACTGACGCCCCT |
| *ACT1* qPCR | Fw | 20 | TCTGAGGTTGCTGCTTTGGT |
| *ACT1* qPCR | Rv | 21 | CCGACGATAGATGGGAAGACA |

**Supplementary Table S3.** List of plasmids used and generated in this work.

| **Plasmid** | **Construct** | **Source** |
|---|---|---|
| pRS426-$P_{GAL1}$-*Luc* | KanMxRV-$P_{GAL1}$-*Luc*-*CYC1*$_{ter}$ | Salinas et al., 2018 |
| pRS426-$P_{5XGAL1}$-*Luc* | KanMxRV-$P_{5XGAL1}$-*Luc*-*CYC1*$_{ter}$ | Salinas et al., 2018 |
| pRS316-FUN-LOV$^{SP\text{-}Hph}$ | HphMxRV-$P_{PGK1}$-WC-1-*GAL4* DBD-*ADH1*$_{ter}$ – $P_{TDH3}$-VVD-*GAL4* AD-*CYC1*$_{ter}$ | Figueroa et al., 2022 |
| pRS316-$P_{GAL1}$-*sfGFP* | KanMxRV-$P_{GAL1}$-*sfGFP*-*CYC1*$_{ter}$ | This work |
| pRS316-$P_{5XGAL1}$-*sfGFP* | KanMxRV-$P_{5XGAL1}$-*sfGFP*-*CYC1*$_{ter}$ | This work |
